# Emergent dynamics of a three-node regulatory network explain phenotypic switching and heterogeneity: a case study of Th1/Th2/Th17 cell differentiation

**DOI:** 10.1101/2021.11.03.465892

**Authors:** Atchuta Srinivas Duddu, Sauma Suvra Majumdar, Sarthak Sahoo, Siddharth Jhunjhunwala, Mohit Kumar Jolly

**Author notes:** Author to whom correspondence to be addressed (M.K.J.).

## Abstract

Naïve helper (CD4+) T-cells can differentiate into distinct functional subsets including Th1, Th2, and Th17 phenotypes. Each of these phenotypes has a ‘master regulator’ – T-bet (Th1), GATA3 (Th2) and RORγT (Th17) – that inhibits the other two master regulators. Such mutual repression among them at a transcriptional level can enable multistability, giving rise to six experimentally observed phenotypes – Th1, Th2, Th17, hybrid Th/Th2, hybrid Th2/Th17 and hybrid Th1/Th17. However, the dynamics of switching among these phenotypes, particularly in the case of epigenetic influence, remains unclear. Here, through mathematical modeling, we investigated the coupled transcription-epigenetic dynamics in a three-node mutually repressing network to elucidate how epigenetic changes mediated by any ‘master regulator’ can influence the transition rates among different cellular phenotypes. We show that the degree of plasticity exhibited by one phenotype depends on relative strength and duration of mutual epigenetic repression mediated among the master regulators in a three-node network. Further, our model predictions can offer putative mechanisms underlying relatively higher plasticity of Th17 phenotype as observed *in vitro* and *in vivo*. Together, our modeling framework characterizes phenotypic plasticity and heterogeneity as an outcome of emergent dynamics of a three-node regulatory network, such as the one mediated by T-bet/GATA3/RORγT.

## Introduction

Differentiation of naïve CD4+ T-cells into diverse T-helper (Th) cells facilitates versatile and adaptable immune responses to different challenges, and serves as a powerful model system to investigate cell-fate decision-making (Evans and Jenner, 2013). Different Th cells – Th1, Th2, Th17 among others – have distinct cytokine and functional profiles. Th1 cells mainly produce IFNγ, and mediate host defense against intracellular bacteria and viruses, while Th2 cells produce IL-4 and are implicated in allergic immune responses. Th17 cells secrete IL-17A, IL-17F and GM-CSF, and act against bacterial and fungal pathogens such as *Mycobacterium* (Kaiko *et al*., 2008; Stadhouders *et al*., 2018). Earlier thought to be mutually exclusive and (terminally) stable phenotypes, recent single-cell evidence has revealed the heterogeneity and plasticity of Th cell subsets. For instance, hybrid Th1/Th2, Th2/Th17 and Th1/Th17 phenotypes have been observed at a single-cell level *in vitro* and *in vivo* (Peine *et al*., 2013; Chatterjee *et al*., 2018; Xhangholi *et al*., 2019; Tortola *et al*., 2020). Moreover, *in vitro* restimulation can drive switching among multiple phenotypes – Th1, Th2, Th17 and the corresponding hybrid ones (Curtis *et al*., 2010; Evans and Jenner, 2013; Tortola *et al*., 2020; Cerboni *et al*., 2021). However, a dynamical characterization of phenotypic switching among these Th cell subpopulations has not been performed.

T-bet, GATA3 and RORγT have been proposed as the ‘master regulators’ of Th1, Th2 and Th17 cells respectively. They can mutually repress each other and self-activate directly or indirectly, thus driving CD4+ naïve cell differentiation into diverse Th subsets (Fang and Zhu, 2017). Such mutual repression is a hallmark of ‘sibling’ cell-fates in many systems, such as for PU.1/GATA1 in the case of common myeloid progenitor differentiating to a myeloid or erythroid fate (Zhou and Huang, 2011), or ZEB1/ GRHL2 for epithelial-mesenchymal transition (Hari *et al*., 2020). Similar to PU.1/GATA1 and ZEB1/ GRHL2, the T-bet/GATA3/RORγT regulatory network can be multistable, enabling the co-existence of different phenotypes and switching among them – Th1 (high T-bet, low GATA3, low RORγT), Th2 (low T-bet, high GATA3, low RORγT), Th17 (low T-bet, low GATA3, high RORγT), hybrid Th1/Th2 (high T-bet, high GATA3, low RORγT), hybrid Th2/Th17 (low T-bet, high GATA3, high RORγT) and hybrid Th1/Th17 (high T-bet, low GATA3, high RORγT). Intriguingly, self-activation of ‘master regulators’ can enrich for hybrid phenotypes (Duddu *et al*., 2020). However, a comprehensive analysis of the interplay among factors influencing the rates of transition among these phenotypes remains to be done.

Epigenetic changes, including modifications of histones and DNA methylation status, can control the rate of phenotypic switching by influencing the access of ‘master regulators’ to their genomewide targets. Thus, cell-specific chromatin landscape and ‘histone code’ can form stable epigenetic marks at various gene loci, thus governing the commitment, heritability and plasticity of various cell-fates (Chang and Aune, 2007; Miyamoto *et al*., 2015; Suelves *et al*., 2016). Early biochemical evidence for the importance of epigenetic processes in T-cell differentiation came from studies showing that treatment of T cells with inhibitors of histone deacetylases (HDACs) or DNA methylation led to the production of IL-2 and IFNγ by cells that could not previously produce them (Wilson *et al*., 2009). Further, T-bet, GATA3 and RORγT have been shown to control chromatin accessibility required for T-cell differentiation into different lineages/phenotypes (Sanders, 2006; Hirahara *et al*., 2011). Interestingly, such influence was shown to be at least partly independent of signals from cytokine receptors/signaling, suggesting many parallel paths that these ‘master regulators’ can take to suppress other lineages and to promote theirs (Josefowicz, 2013; Lee *et al*., 2020). Epigenetic changes such as chromatin remodeling are crucial not only for CD4+ T-cell differentiation, but also for lineage stability (Renaude *et al*., 2021). For instance, in Th2 cells, repressive histone marks H3K27me3 and H3K9me3 are deposited on Th1-associated genes, thus forming hetero-chromatin at those specific promoters and establishing transcriptional silencing. Such epigenetic changes drive lineage stability and often need to be ‘erased’ for reprogramming.

Once differentiated, T-helper subsets are often reprogrammed to other ones during injury response and resolution (Tortola *et al*., 2020). The stronger the lineage stability mechanisms in a cell, the less reprogrammable it can become. While mutual antagonism at transcriptional and epigenetic levels enable the establishment of a cell-state and have been witnessed in other cellular decisionmaking contexts (Tripathi *et al*., 2020; Serresi *et al*., 2021), they can differently control the reprogramming rates. For instance, in various instances, ‘epigenetically locked’ cells can be difficult to reprogram (Nashun *et al*., 2015; Baumann *et al*., 2019; Eichelberger *et al*., 2020). Despite such wealth of molecular and functional data for T-cell differentiation, we still lack a quantitative systems-level investigation how the rates of phenotypic switching among Th1, Th2 and Th17 depend on the epigenetic influences mediated by T-bet, GATA3 and RORγT on each other.

Here, we build on our previous study that showed that a mutually repressing three-node system enables three pre-dominant states driven by each node, and that switching between these states was possible. However, the dynamics of switching and longevity (mean residence times) of these phenotypes, especially in cases where epigenetic control exists, remained unexplored. While Th cells are known to undergo similar switching, experimental data regarding the dynamics of the switches is scarce and the experiments are difficult to perform. Thus, we have used a mathematical modeling approach to address the questions of emergent dynamics of a three-node system repressing each other at both transcriptional and epigenetic levels. Our simulations reveal that the rate of switching among phenotypes and consequent changes in population distribution of cellular phenotypes is a function of the relative strength as well as duration corresponding to epigenetic repression mediated by the three ‘master regulators’ on one another. The stronger the incoming epigenetic repression for a given master regulator, the higher the probability of switching out of the corresponding phenotype. These results unravel a potential design principle of T-cell differentiation at individual cell and population levels.

## Results

### Transcriptional dynamics of Th1, Th2 and Th17 induction and corresponding phenotypic plasticity

To assess the transcriptional dynamics of T-cell differentiation, we quantified the extent of changes in Th1, Th2 and Th17 signature gene-sets under diverse experimental conditions using publicly available datasets. First, we analyzed a RNA-seq dataset (GSE71645) in which naïve T-cells were cultured for 72 hours either in a Th1 inducing medium (IL-12 treatment) or in a Th2 inducing medium (IL-4 treatment) (Kanduri *et al*., 2015), and projected the samples on a two-dimensional plane of Th1 and Th2 ssGSEA (single-sample Gene Set Enrichment Analysis) scores (Subramanian *et al*., 2005) (**Fig 1A, i**). We observed that cells treated with IL-12 showed a significant enrichment in Th1 signature while IL-4 treated cells showed an enrichment in Th2 signature (**Fig 1A, ii**). Time-course microarray data collected for this experiment (GSE71566) demonstrated that relative enrichment of one of the two signatures (Th1 or Th2) can be seen as early as three days in culture (**Fig 1B**). Similar mutually opposing trends for Th1 and Th2 enrichments were also seen in another independent dataset (GSE62484) that contained populations of naïve T cells, activated Th1 and activated Th2 cells (**Fig 1C**) (Hertweck et al., 2016). Analysis of two other time-course transcriptomic datasets (GSE60678, GSE32959) reinforced our observations that a three day treatment of naïve CD4+ T with either a Th1 or a Th2 inducing medium began to show differential activation of Th1 or Th2 induction programs (Äijö *et al*., 2012; Gustafsson *et al*., 2015), and stabilized at later time points (**Fig 1D, 1E**). These dynamics are reminiscent of many cell differentiation trajectories where a multi-potent progenitor cell often co-expresses mutually opposing ‘master regulators’ (and/or their targets) corresponding to two (or more) phenotypes. This multi-potent state of a cell is destabilized under the impact of exogenous signals (cytokines, growth factors etc.) that push it towards an “attractor” corresponding to one of the differentiated states (Huang *et al*., 2007; Bargaje *et al*., 2017).

**Fig 1:**
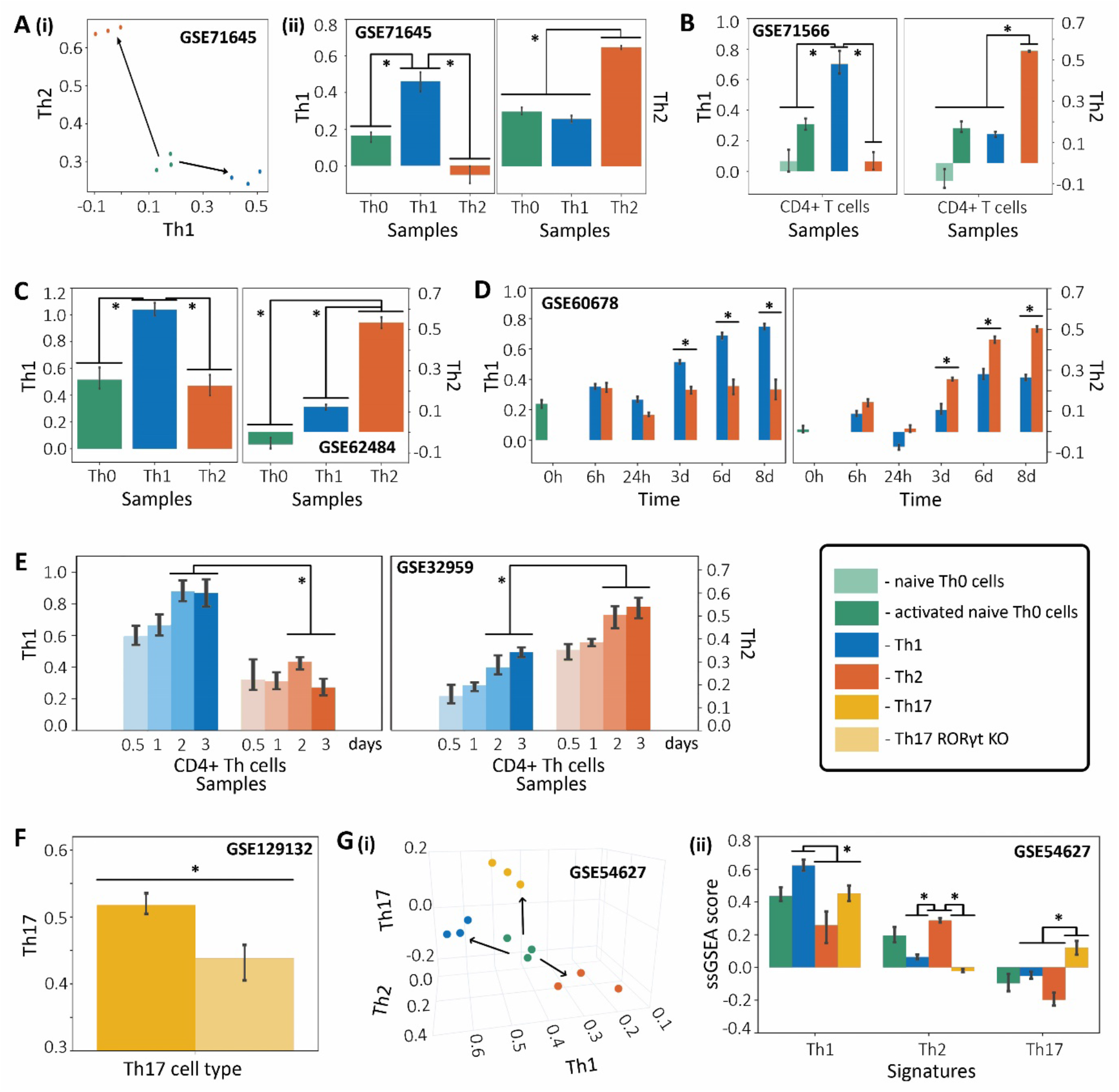
Transcriptomic analysis showing enrichment of Th1, Th2 and Th17 signatures specific to corresponding cell types. **A) i)** 2D scatterplot showing T naïve, Th1 and Th2 cell types on the Th1-Th2 ssGSEA score plane. **ii)** Quantification of differences in levels of Th1 and Th2 ssGSEA scores across T naïve, Th1 and Th2 cell types (GSE71645). **B)** Quantification of differences in levels of Th1 and Th2 ssGSEA scores across T naïve, activated T naïve, induced Th1 and induced Th2 cell types (GSE71566). **C)** Quantification of differences in levels of Th1 and Th2 ssGSEA scores across T naïve, activated Th1 and activated Th2 cell types (GSE62484). **D)** Quantification of differences in levels of Th1 and Th2 ssGSEA scores across T naïve, Th1 and Th2 cell types over the timepoints 0 hours, 6 hours, 1 day, 3 days, 6 days, and 8 days (GSE60678). **E)** Quantification of Th1 and TH2 ssGSEA scores for differentiating Th1 and Th2 cells over time points 0.5, 1, 2, and 3 days (GSE32959). **F)** Quantification of differences in levels of Th17 ssGSEA scores across WT and RORγT knockout Th17 cells (GSE129132). **G) i)** 3D scatterplot showing T naïve, Th1, Th2 and Th17 cell types on the Th1-Th2-Th17 ssGSEA score space in a non-treatment condition (control set at 0 hours) and **ii)** its corresponding (same condition) quantification of differences in the levels of Th1, Th2 and Th17 signatures (ssGSEA scores) (GSE54627). (* denotes a significantly different level of ssGSEA scores assessed by Students t-test; p-value < 0.05).

Next, we investigated transcriptomic signatures corresponding to Th17 differentiation, and noticed that this signature captured the inhibition of Th17 differentiation when RORγT, a ‘master regulator’ of Th17 cell state, was silenced in naïve CD4+ T-cells (**Fig 1F**) (GSE123192) (Lee *et al*., 2020). Finally, we assessed how the three different cell types (Th1, Th2, Th17) are separated in a threedimensional space of their ssGSEA scores (**Fig 1G**). We found that each of the cell types displayed significant enrichment of their corresponding signatures. Further, the T naïve cells are situated “intermediate” to the three cell type signatures (**Fig 1G, i**), consistent with the undifferentiated state co-expressing markers of multiple phenotypes it can give rise to, as seen across biological contexts (Olsson *et al*., 2016). We observed that Th1, Th2 and Th17 cells showed significant enrichment in their respective signatures while the signatures of the competing programs (for example, Th1 and Th2 programs during Th17 differentiation) were significantly suppressed (**Fig 1G, ii**). Collectively, these results indicate the robust transcriptomic signatures associated with Th1, Th2 and Th17 induction during the trifurcation event during T cell differentiation.

T-bet, GATA3 and RORγT – the proposed ‘master regulators’ of Th1, Th2 and Th17 respectively – and their targets constitute the above-mentioned transcriptomic signatures associated with Th1, Th2 and Th17 differentiation (Radens *et al*., 2020). They have been reported to mutually repress each other, thus pushing CD4+ naïve cell into diverse differentiation trajectories (Fang and Zhu, 2017), a trend robustly captured in transcriptomic datasets shown above. Thus, a network of three mutually repressing regulators (A, B and C) can serve as a model for CD4+ T-cell differentiation.

We have previously shown that such a ‘toggle triad’ among A, B and C (**Fig 2A**) can enable three predominant states – (high A, low B, low C), (low A, high B, low C) and (low A, low B, high C) (**Fig 2B**) (represented by Abc, aBc and abC correspondingly hereafter) (Duddu *et al*., 2020). The states enabled by ‘toggle triad’ are reminiscent of emergent dynamics of ‘toggle switch’ – a mutually inhibitory feedback loop between two master regulators that often enable two mutually exclusive states: (high A, low B) and (low A, high B), corresponding to a specific phenotype (Cherry and Adler, 2000; Gardner *et al*., 2000; Graham *et al*., 2010). A toggle switch explains the behavior of a progenitor cell differentiating into one of two cell fates, each fate driven majorly by a master transcription factor (TF). Similar to phenotypic plasticity and heterogeneity observed in a toggle switch under the influence of noise (Gardner *et al*., 2000; Ozbudak *et al*., 2004), we would expect the three states enabled by a ‘toggle triad’ also to be capable of switching among one another. To confirm this, we performed stochastic switching simulations for six different tristable parameter sets (P1-P6) that revealed possible switching among the three phenotypes - Abc (blue), aBc (red) and abC (yellow) (**Fig 2C**). The mean residence times in each state varies with the parameter set. However, across parameter sets, none of the states could be classified as transient/intermediate. For further characterization of the dynamical traits of the toggle triad, we mapped two phase diagrams for a representative parameter set with degradation or production rates of B and C (kB, kC; gB, gC) as the respective bifurcation parameters (**Fig S1**).

**Fig 2:**
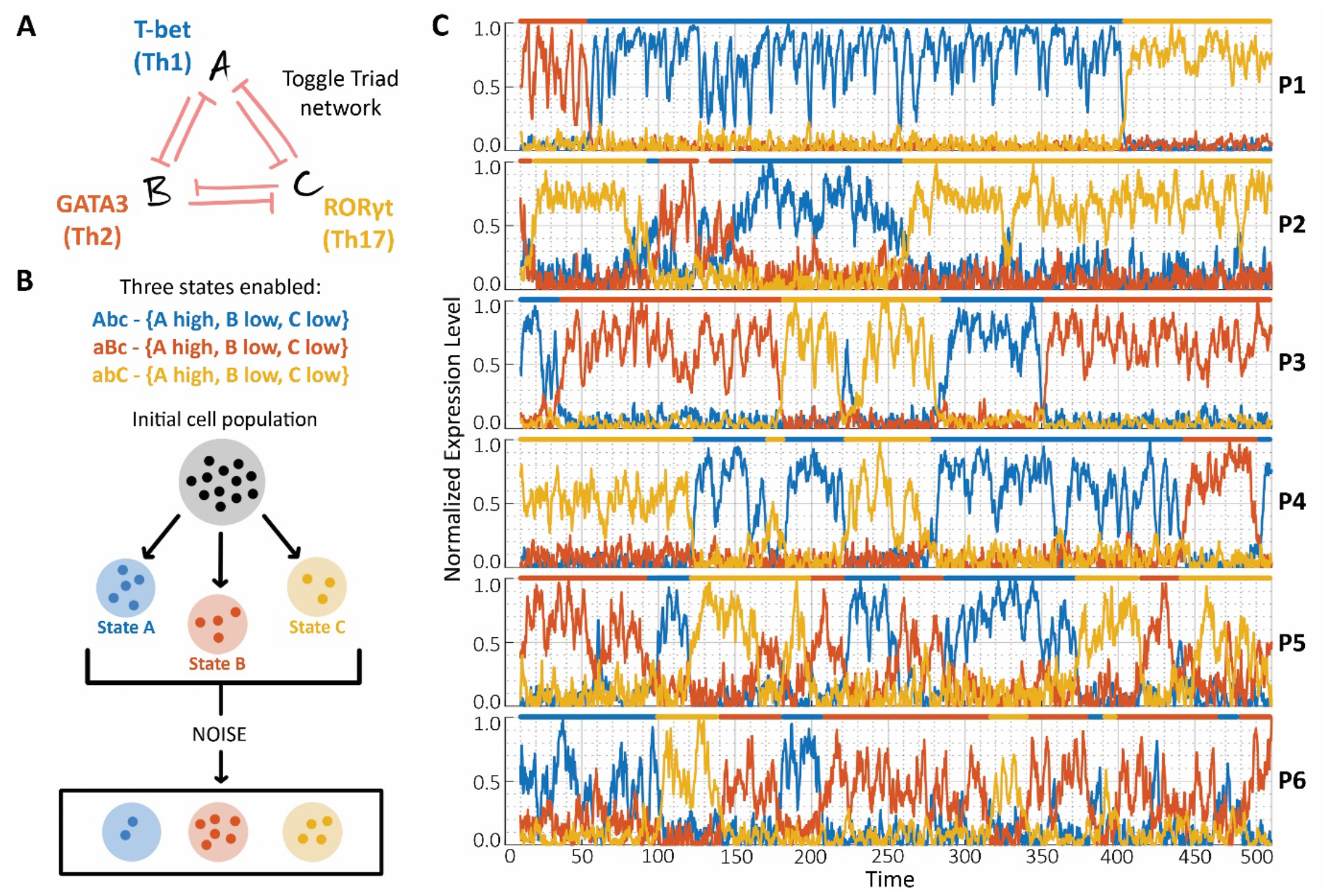
Phenotypic heterogeneity in T-helper cell population. **A)** Toggle Triad network topology underlies the differentiation of naïve CD4+ T-cell into T-helper cells (Th1, Th2 and Th17). Each master regulator (T-bet, GATA3 and RORγT respectively) mutually represses the other two. **B)** The three states enabled by a toggle triad are listed along with notation used hereafter. A schematic representing a kinetic model with certain parameter set enabling a heterogenous population and noise enabling switching between the states. **C)** Stochastic simulations of the network for different parameter sets (P1-P6 (Table S1)) showing switching between the three states.

Together, these results indicate that depending on the relative abundance of T-bet, GATA3 and RORγT, cells can exist in one or more of the three dominant phenotypes (Th1, Th2 and Th17) and can switch back and forth under the influence of stochastic fluctuations (biological noise). Such state-switching can induce and maintain phenotypic heterogeneity in a given T-helper cell population, with the relative frequencies of Th1, Th2 and Th17 dependent on relative levels of the master regulators (or equivalently, the concentration of different cytokines which can drive various cell-fates through their action on these master regulators).

### Epigenetic repression driven by a master regulator can enrich for its corresponding phenotype in a heterogeneous population

Besides mutual repression at a transcriptional level, the three ‘master regulators’ (T-bet, GATA3 and RORγT) can engage in epigenetic mutual repression as well (Mukasa *et al*., 2010; Wei *et al*., 2010; Zhu *et al*., 2012; Sasaki *et al*., 2013; Lee *et al*., 2020). To incorporate epigenetic repression in our framework that captures transcriptional repression among these three ‘master regulators’, we utilized a phenomenological model approach (Miyamoto *et al*., 2015) that introduces an epigenetic parameter (α) to quantify the threshold (half-maximal) levels corresponding to the influence of expression levels of one node on its target. The higher the value of α corresponding to a network edge, the stronger is the epigenetic repression incorporated in that interaction. The underlying idea behind this framework is that epigenetic remodeling (or repression) serves as a self-stabilizing mechanism to maintain a cell-state and potentially propagate it across generations. In this epigenetic modeling framework, the longer a node stays at a high expression level (“ON”), the less likely it is for the node to switch to a lower expression level (“OFF”, due to chromatin and/or DNA methylation changes it may have mediated meanwhile), or in other words, the more likely a cell is to maintain the state driven by that ‘master regulator’, even if the levels of that node decline later. For instance, in fully matured Th1 cells, IFN-γ expression becomes relatively independent of T-bet activity and coincides with DNA methylation changes (Mullen *et al*., 2002).

Here, we use this framework to simulate multiple scenarios, i.e. epigenetic repression incorporated on various edges in a network, and quantify changes in phenotypic distribution in a differentiating T-cell population (Th1, Th2, Th17), in presence of noise to account for stochastic effects. First, we considered the scenario of epigenetic repression mediated by one of the master regulators (say, B) on inhibitory links to other two nodes (from B to A, and from B to C) (**Fig 3A**). The population distribution is calculated by considering multiple initial conditions (here, 1000), each representing an individual cell. The trajectory of each initial condition (cell) is followed, and the expression values of A, B and C are noted. Depending on the expression levels of the nodes, the state (of the cell) is defined and the population distribution is deduced. Noise is incorporated into the system at definite time intervals (See Methods). For a given tristable parameter set, we first identified population distribution in absence of any epigenetic repression (α_BA_ = 0, α_BC_ = 0) – approximately 20% cells were present in (high A, low B, low C) (Abc or state A) and 30% in (low A, high B, low C) (aBc or state B) states, while 50% cells exhibited a (low A, low B, high C) (abC or state C) phenotype (**Fig 3B**, left bottom panel).

**Fig 3:**
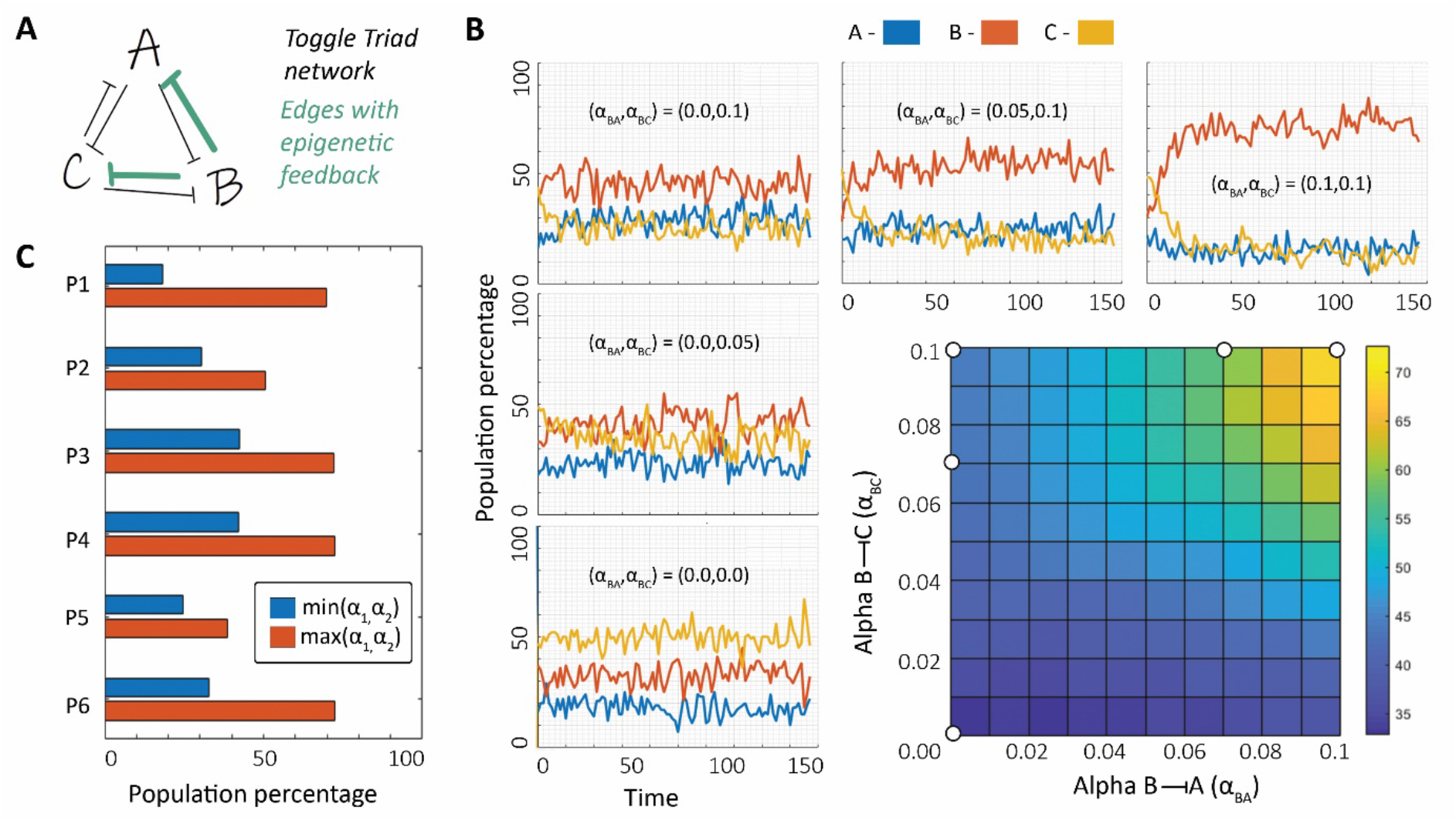
Epigenetic repression mediated by one node in toggle triad on the other two nodes. **A)** Toggle triad network topology in which interactions incorporating epigenetic repression are marked in green. **B)** Phase plot showing the population percentage of cells in state A (Abc) with bifurcation parameters as the α values corresponding to the epigenetic feedback of B -| A and B -| C, as well as dynamics of distribution of population percentage between states A, B and C for certain pairs of α_BA_ and α_BC_ values. **C)** Population percentage of the node from which interactions with epigenetic repression originate, with pair of α values at corresponding maximum and minimum for six parameter sets (P1-P6; given in Table S1). Parameter set P6 used in **B.** Results for parameter sets P1-P5 shown in Fig S2, S3.

As we increase the strength of epigenetic repression from B to C (α_BC_), we observe a decrease in population percentage of abC (or C) state and a corresponding increase in that corresponding to A and B. At (α_BA_ = 0, α_BC_ = 0.05), the system displays a population percentage distribution of about 25% A, 45% B and 30% C. This trend continues with further increase in value of α_BC_; at (α_BA_ = 0, α_BC_ = 0.1), a population distribution around 30% A, 45% B and 25% C is observed (**Fig 3B**, left column). Incorporating epigenetic repression from B to A (α_BA_) in addition to α_BC_ = 0.1 further increases the population percentage corresponding to B. At (α_BA_ = 0.1, α_BC_ = 0.1), the population predominantly consists of cells in state B (70%) (**Fig 3B**, top row). This trend is further exemplified by the phase plot showing that the population percentage of B is minimum at low values of (α_BA_, α_BC_) and increases sharply as (α_BA_, α_BC_) values increase (**Fig 3B**). We performed similar analysis for other tristable parameter sets and observed similar trends, although the degree of enrichment of corresponding state varied (**Fig 3C, S2, S3**). For instance, in parameter sets P1, P3 and P4, including such epigenetic repression drastically alters the phenotypic distribution in favor of the master regulator which is inhibiting the other two ones epigenetically (C inhibits A and B in P1, in B inhibits A and C in P3, B inhibits A and C in P4). The trends are consistent but not as strong, however, in parameter sets P2 (A inhibits B and C) and P5 (C inhibits A and B) (**Fig S2, S3**).

Put together, we conclude that including epigenetic repression from interactions originating from one of the three nodes in a toggle triad help increase the percentage of cells in a state for which that node serves as a ‘master regulator’. The magnitude of change in phenotypic composition depends on the strength of epigenetic feedback of either or both interactions. Extrapolating these results in the context of T-cell differentiation, they imply that if T-bet can epigenetically repress on GATA3 and/or RORγT, the population predominantly will consist of Th1 cells. Similarly, Th2 (or Th17) cells can be the predominant phenotype in a heterogeneous T-cell population if GATA3 (or RORγT) can repress T-bet and RORγT (or T-bet and GATA3) epigenetically.

These observations offer dynamical insights into how the ability of RORγT to “not only establishing the permissive epigenetic landscape but also preventing the repressive one” (Lee *et al*., 2020) is crucial for robust Th17 differentiation. Th17 cells have been reported to be relatively more plastic, and can be reprogrammed readily to Th1 and Th2 (Lexberg *et al*., 2008; Stadhouders *et al*., 2018; Cerboni *et al*., 2021), with implications in diseases such as rheumatoid arthritis (Yang *et al*., 2019). This instability has been suggested to be driven by rapid epigenetic modifications for cytokines (*Il17a-Il17f, Ifng*) and transcription factor (*Rorc*) gene expression associated with Th17 cell lineage specification (Mukasa *et al*., 2010). Our simulations proposed that increased plasticity of Th17 cells may be a consequence of a) weak epigenetic repression driven by RORγT on T-bet and/or GATA3, and/or b) strong epigenetic repression mediated by T-bet and/or GATA3 on RORγT.

### Impact of competing and complementing epigenetic repression driven by two master regulators on the population distributions

Next, we consider the scenario of epigenetic repression incorporated on a pair of mutual repressive links (marked by green in **Fig 4A**: here, inhibition between B and C is considered). The strength of epigenetic repression is characterized by corresponding α values α_BC_ and α_CB_. For the given parameter set (P6), the system converges to approximately 20% A, 30% B and 50% C in absence of any epigenetic repression (α_BC_ = 0, α_CB_ = 0) (**Fig 4A**). Increasing either α_BC_ or α_CB_ increases the population percentage corresponding to state B or C respectively: at (α_BC_ = 0.2, α_CB_ = 0), population distribution is about 30% A, 45% B and 25% C, while at (α_BC_ = 0, α_CB_ = 0.3), the heterogeneous population comprises of 15% A, 15% B and 70% C. Increasing both α_BC_ and α_CB_ values brings the population closer to the case of no epigenetic influence; at (α_BC_ = 0.2, α_CB_ = 0.3), the population distribution comprises of approximately 25% A, 25% B and 50% C. Quantifying the ratio of population percentages corresponding to states B and C with α_BC_ and α_CB_ as the two parameters, we observed skewed ratios of the two phenotypes, when one of the epigenetic repression links is much stronger than the other (< 1 at α_BC_ = 0.2, α_CB_ = 0; and > 6 at α_BC_ = 0, α_CB_ = 0.3) (**Fig 4B**). Similar analysis for other parameter sets (**Fig 4C, Fig S4-S8**) substantiates these trends, where the ratio of population percentages of two representative states is skewed when one of the links dominates ((α_1–min_, α_2-max_) and (α_1-max_, α_2-min_)) but not when the epigenetic influence on one another is of comparable strengths ((α_1-min_, α_2-min_) and (α_1-max_, α_2-max_)). In the context of T-cell differentiation, these results imply that if T-bet and GATA3 can repress the expression or function of each other at an epigenetic level, the relative strength of their epigenetic inhibitions governs the relative proportions of Th1 and Th2 in a heterogeneous population. Similar statements can be made for mutual repression between any other pair of ‘master regulators’ here.

**Fig 4:**
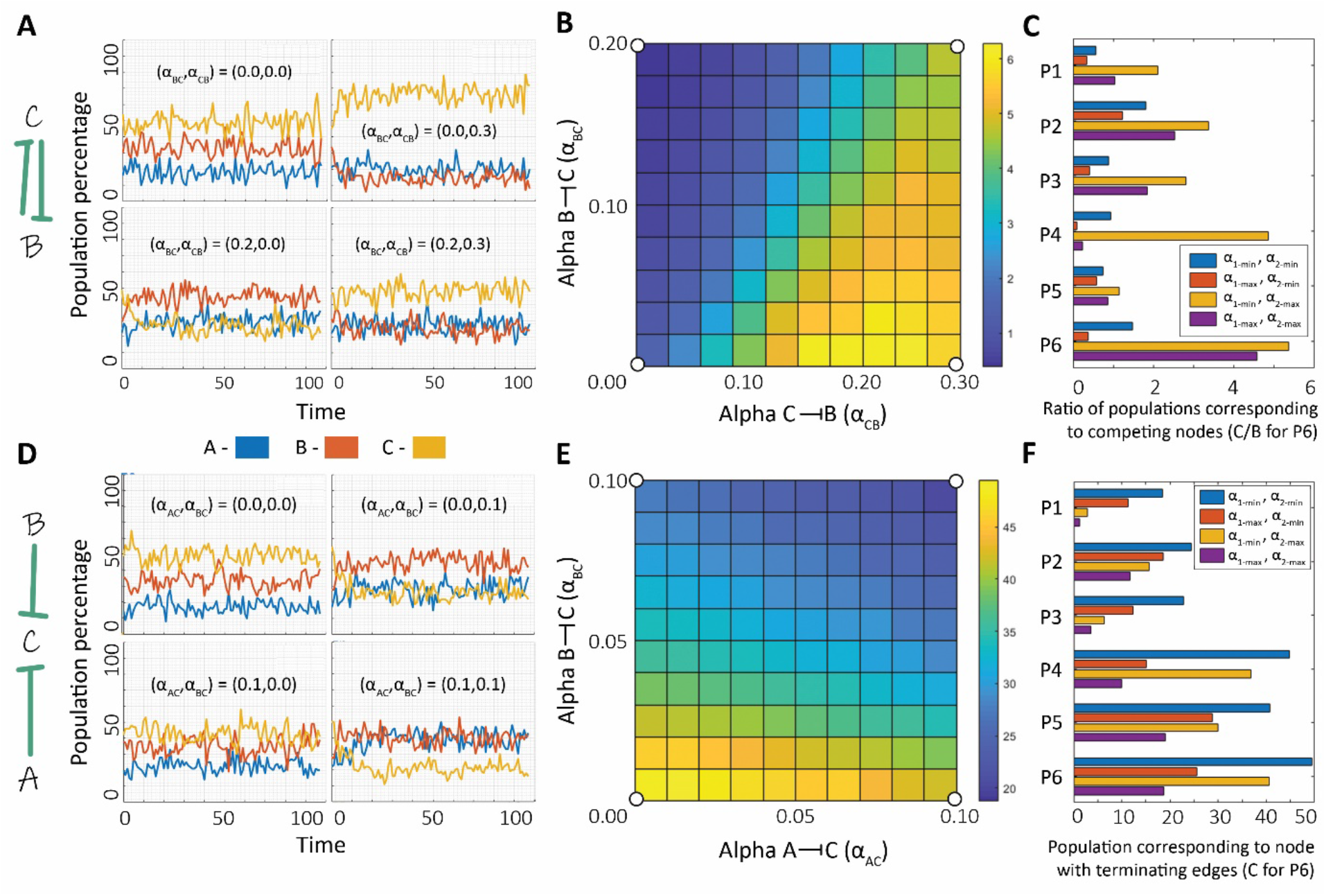
Epigenetic repression on edges originating from more than one node in the toggle triad. **A)** (Left) Toggle triad network in which interactions where epigenetic repression is incorporated are marked in green. (Right) Dynamics of distribution of population percentage between states A, B and C for certain pairs of α_BC_ and α_CB_ values. **B)** Phase plot showing the ratio of population percentage of C to that of B with bifurcation parameters as the α values corresponding to the epigenetic feedback of B -| C and C -| B. **C)** Ratio of population percentages of the nodes from which interactions with epigenetic feedback originate with pair of α values at combinations of maximum and minimum for three different parameter sets. **D)** (left) same as A). (right) same as A) but for certain values of α_AC_ and α_BC_. **E)** Phase plot showing the population percentage of C with bifurcation parameters as the α values corresponding to epigenetic feedback of A -| C and B -| C. **F)** Population percentage of C (for parameter set P6), that of corresponding nodes in other parameter sets (P1-P5). Results for P6 are shown in panels B and E; those for P1-P5 shown in Fig S4-S8.

Finally, we considered a scenario where both epigenetic repression is incorporated on two edges terminating at a single node of the toggle triad (marked in green in **Fig 4D**; here, inhibition of C by A and by B). The strength of the epigenetic feedback is characterized by corresponding α values, α_AC_ and α_BC_ respectively. Without any epigenetic feedback (α_AC_ = 0, α_BC_ = 0), the system equilibrates to a population percentage distribution of 20% A, 30% B and 50% C (**Fig 4D**). Increasing either α_AC_ or α_BC_ decreases the population percentage corresponding to state C (abC); at (α_AC_ = 0.1, α_BC_ = 0) and (α_AC_ = 0, α_BC_ = 0.1), the population percentage corresponding to state C drops to about 40% and 25% respectively. Increasing both α_AC_ and α_BC_ further reduces the population percentage of state C (20% C at (α_AC_ = 0.1, α_BC_ = 0.1)) (**Fig 4D, E**). Similar trends are seen for other parameter sets where one node is epigenetically being repressed by other two nodes: A inhibited by B and C epigenetically; C inhibited by A and B epigenetically; B inhibited by A and C epigenetically) (**Fig 4F, S4-S8**).

Together, these three different scenarios underscore how epigenetic repression incorporated through different inhibitory edges in T-bet/GATA3/RORγT regulatory network can alter the population distribution structure (% of Th1, Th2, Th17) in a T-cell differentiation context.

### The impact of epigenetic influence on population distribution depends both on corresponding strength(s) and duration(s)

So far, we have considered epigenetic feedback on any edge to not vary as a function of time. Next, we characterize the dynamics of the system with the epigenetic feedback provided only for certain time duration instead of being present constantly (throughout the simulation) as previously.

We considered the tristable parameter set P6, where epigenetic repression was incorporated on two interactions originating from a single node (**Fig 5A**; B inhibiting both A and C; strengths: α_BA_ and α_BC_, same as the case considered in **Fig 3**). We switched on the epigenetic feedback for only a fraction of entire simulation time (X). Without any epigenetic feedback, the population distribution converged to about 32% in states A and B, and 36% in state C (**Fig 5B**). As X increased, the population percentage corresponding to state B increases while those corresponding to A and C simultaneously decrease. We then varied both parameters – X and α_BA_ (= α_BC_) - to make a phase plot. The population percentage corresponding to B is about 33% at α_BA_ = α_BC_ = 0 and no epigenetic feedback (X = 0). At low strengths (α_BA_ = α_BC_ < 0.05) and short durations (X < 0.5) of epigenetic feedback, the population distribution remains largely unperturbed (left bottom of **Fig 5C**). However, beyond this approximate threshold, increasing either the strength (dose) of epigenetic influence or the duration (marked by an asterisk and arrows in **Fig 5C**), leads to significant changes in the population levels corresponding to B.

**Fig 5:**
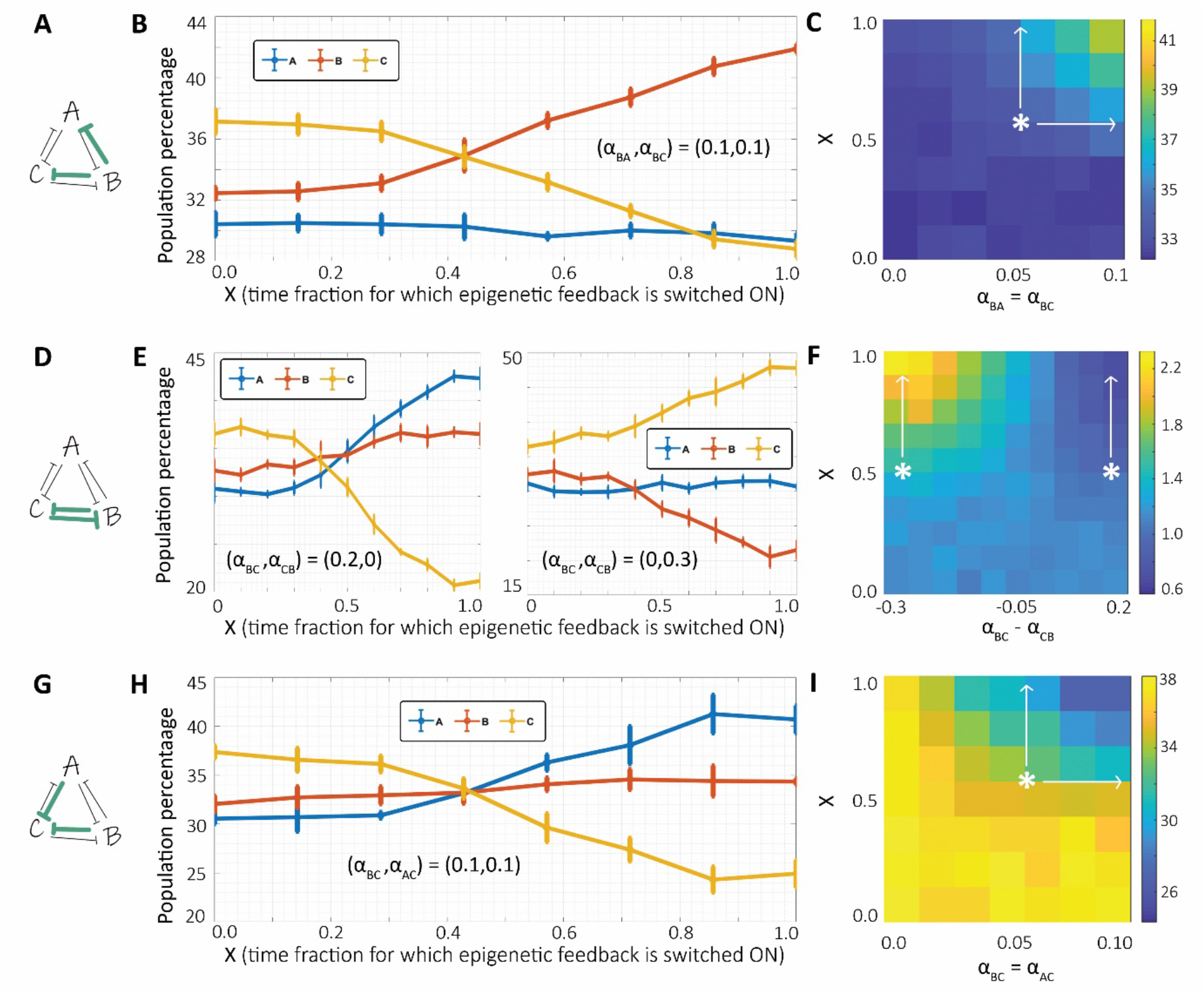
Epigenetic repression on edges originating from more than one node in the toggle triad. **A)** Toggle Triad network topology in which interactions marked in green are being provided with epigenetic feedback. **B)** Population percentages of A, B and C as X, the fraction of time for which the epigenetic feedback for both marked interactions is switched ON and then turned OFF. **C)** Phase plot showing population percentage of B (node from which interactions with epigenetic feedback originate) with bifurcation parameters as the α value corresponding to epigenetic feedback of B -| A and B -| C (α_BA_ = α_BC_) and X. **D)** Same as A) **E)** Same as B) but for two cases where feedback for one of the interactions, B -| C (C -| B) is switched ON with the other one C-|B (B-|C) switched OFF. **F)** Phase plot showing ratio of population percentage of C to B (nodes between which interactions with epigenetic feedback are present) with bifurcation parameters as the difference of α values corresponding to the epigenetic feedback of B -| C and C -| B and X. **G)** Same as A). **H)** Same as B) **I)** Phase plot showing population percentage of C (node onto which interactions with epigenetic feedback terminate) with bifurcation parameters as the α value corresponding to the epigenetic feedback of A-|C and B-|C (both same so considered on single axis) and X. Results for parameter set P6 shown here; those for P1-P5 shown in Fig S9-S13.

Next, we considered the case where epigenetic influence was incorporated for mutual inhibition between two nodes (**Fig 5D**; B inhibiting C and C inhibiting B; strengths: α_BC_ and α_CB_; same as the case considered in **Fig 4A-B**). Without any epigenetic feedback, the population distribution converged to about 32% in both the states A and B, and 36% in state C (**Fig 5B**). We started with case where one of the two ‘master regulators’ inhibited the other one epigenetically (B inhibits C: α_BC_ = 0.2, α_CB_ = 0) and increased X (**Fig 5E**, left). The population percentage corresponding to C decreased (from about 37% to 20%) while that corresponding to B increased (from about 32% to 37%), although not drastically. We then considered the scenario of epigenetic repression through the other interaction (i.e., C inhibits B epigenetically - α_BC_ = 0, α_CB_ = 0.3), and estimated populations distributions at varying values of X (**Fig 5E**, right). As X increased, the population percentage corresponding to B dropped sharply (from about 32% to 20%) and that corresponding to C increased concurrently (from about 37% to 47%).

Next, to evaluate the impact of mutual epigenetic repression more clearly, we plotted a phase diagram by varying two parameters – X and the difference between epigenetic repressions that B and C have on each other (=α_BC_ – α_CB_) (**Fig 5F**) showing the ratio of population percentages corresponding to states B and C. When both B and C inhibit each other comparably (|α_BC_ - α_CB_| < 0.05), the population ratio remains largely unchanged irrespective of the duration of feedback X. On the other hand, when either interaction is much stronger than the other (|α_BC_ - α_CB_| >=0.05), the ratio of populations remains largely similar until the duration of epigenetic feedback crosses an approximate threshold (marked by asterisks), after which the population distribution diverges depending on the relative mutual strength of epigenetic influence.

Further, we considered the case with epigenetic repression incorporated on two interactions terminating on the same node (**Fig 5G**; A and B both inhibiting C; strengths: α_AC_ and α_BC_; same as the case shown in **Fig 5D-E**). Without any epigenetic influence, the population converged to about 32% cells in state A and B and 36% in state C (**Fig 5H**). As X is increased, the population percentage corresponding to state C decreases while that corresponding to states A and B concomitantly increase. When we varied both the parameters – X and α_AC_ (= α_BC_) – to draw a phase plot, we found the population percentage corresponding to state C is ~ 38% at α_BC_ = α_AC_ = 0 and no epigenetic feedback (X = 0). At low strength (α_BC_ = α_AC_ < 0.1) and short durations (X < 0.5) corresponding to epigenetic repression, no major changes are observed for the population distribution (left bottom part in **Fig 5I**). However, beyond this approximate threshold (marked by an asterisk in **Fig 5I**), increasing either the strength or the duration (vertical and horizontal arrows in **Fig 5I**) leads to comparable and pronounced decreased in the population corresponding to state C. Similar simulations for the three cases of epigenetic repression considered are performed for other parameter sets (P1-P5) and the trends remain consistent (**Fig S9-S13**).

Put together, we conclude that both the factors – strength of epigenetic silencing (α) or the time duration for which it is switched on (X) – can act independently and alter the population distribution patterns, given a threshold amount of the other. These two variables seem to have additive and complementary effects rather than redundant ones. In terms of T-cell differentiation, these results indicate that either a strong epigenetic silencing of other cell lineages for a short duration or a gradually accumulating impact of epigenetic silencing (DNA methylation, histone modification etc.) can both drive changes in the underlying population heterogeneity, suggesting an ‘area under the curve’ dynamical principle.

Previously, we considered varying values of α_AC_ = α_BC_ and observed how this feedback strength and X affect the phenotypic heterogeneity of the population distribution. Next, we consider a more generic scenario where α_AC_ and α_BC_ values need not be identical. Also, instead of including the epigenetic influence on only incoming links on C (C being inhibited by A and B), we now also incorporate the epigenetic influence of C inhibiting A and/or B, to represent the mutual epigenetic repression scenario.

First, we choose a tristable parameter set ({Abc, aBc, abC}) and include epigenetic influence from C to B with (α_CB_ = 0.2). We can continuously decrease the strength of this influence, i.e. α_CB_ varies between 0 and 0.2, and increase the epigenetic influence from B on C (α_BC_). Thus, similar to mutual repression as seen at a transcriptional level in a toggle switch (Gardner *et al*., 2000), two nodes can also inhibit each other at an epigenetic level as well. The difference between the two parameters (α_BC_ - α_CB_) indicates which epigenetic repression (from B to C or from C to B) is predominant. Thus, we varied two parameters - α_BC_ - α_CB_ and X - and obtained the phase plots corresponding to percentage population in the three states: Abc (state A), aBc (state B) and abC (state C) (**Fig 6A, i-iii** respectively). As α_BC_ - α_CB_ changes from −0.2 to 0.2, i.e. as the epigenetic influence of B inhibiting C takes over that of C inhibiting B, and given a minimal critical value of X (= 0.01), the population corresponding to state B increases from about 12% to about 20%. For the same change in parameters, population for state C decreases correspondingly from about 78% to about 70%. But, for the same change in epigenetic influence at a higher value of X (= 0.03), the change in population corresponding to state B is more drastic (from 12% to 40%), with a correspondingly sharp fall in population corresponding to state C (from 80% to 50%). Similar trends are noted for a different representative example of mutual epigenetic repression is considered (i.e. when A and C inhibit each epigenetically) (**Fig S14A**).

**Fig 6:**
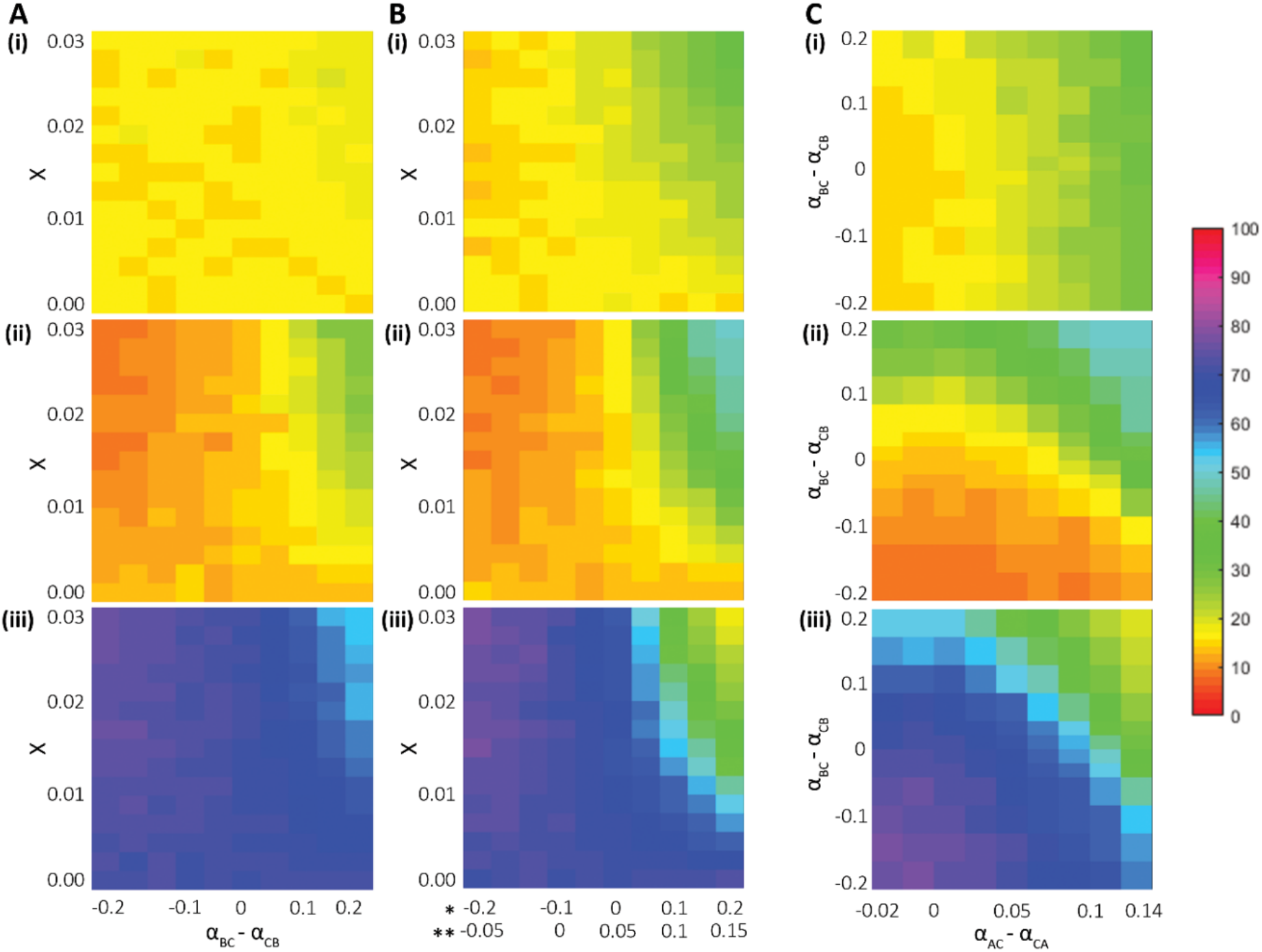
Varying the epigenetic repression from a node to onto the node in toggle triad. **A) i)** Phase plot of population percentage corresponding to node A with variations in two parameters: X and the relative epigenetic influence which is varied from a case of stronger repression from C to B (α_BC_ < α_CB_) to a stronger repression from B to C (α_BC_ > α_CB_) **ii)** Same as i) but for state B. **iii)** Same as i) but for state C. **B) i)** Same as A; i) but for varying relative epigenetic influence in both feedback loops (between A and C, and between B and C). Corresponding α_BC_ - α_CB_ and α_AC_ - α_CA_ values are given on x-axis, marked by * and ** respectively. **ii)** Same as i) but for state B. **iii)** Same as i) but for state C**. C) i)** Phase plot of population percentage corresponding to state A with bifurcation parameters as the difference in α values corresponding to mutual epigenetic repression (α_BC_ - α_CB_ *and* α_AC_ - α_CA_), at X = 0.03. **ii)** Same as i) but for state B. **iii)** Same as i) but for state C. Results for parameter set P6 shown here; those for P1-P5 parameter sets shown in Fig S14-S18.

**Fig 7:**
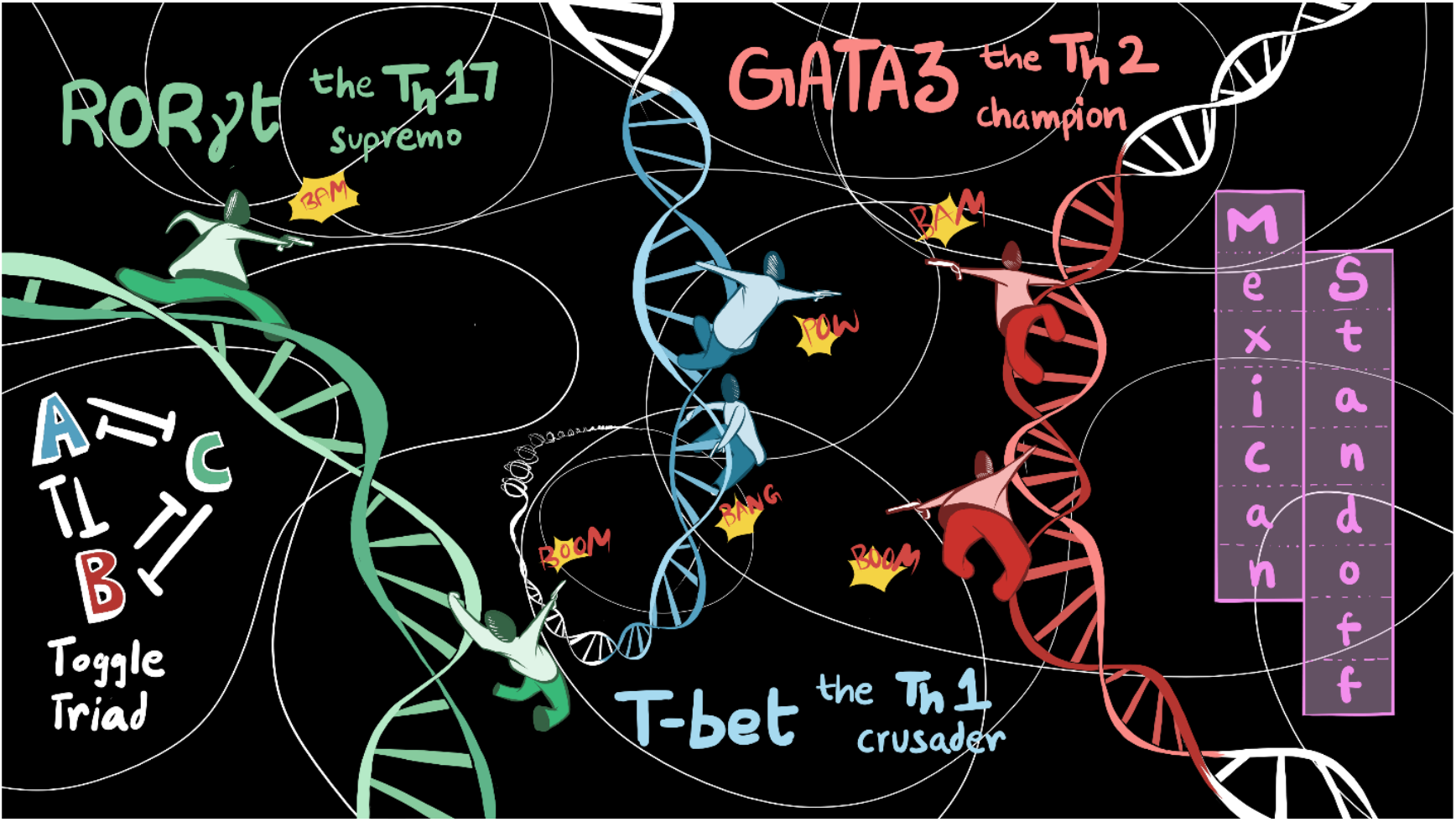
A schematic representing Th1/Th2/Th17 differentiation mediated by a toggle triad.

Next, we investigate the scenario of mutual epigenetic repression for two toggle switches instead of just one: between B and C, as well as between A and C. Thus, in addition to changes in α_BC_ – α_CB_, α_AC_ – α_CA_ can also be considered as a parameter to be varied. For this parameter set, while α_BC_ – α_CB_ changes from −0.2 to 0.2, α_AC_ - α_CA_ changes from −0.05 to 0.15. We observed that at X= 0.03, as the magnitude of the incoming epigenetic repression on C increases as compared to the ability of C to epigenetically repress A and/or B (i.e., α_BC_ - α_CB_ > 0 and α_AC_ - α_CA_ > 0), the population corresponding to state C drops sharply from 80% to 15%, with corresponding increase in population corresponding to both the states A and B (from 10% to 35% for A, and from 10% to 45% for B) (**Fig 6B, i-iii**).

Finally, we consider the case of a fixed duration for which epigenetic repression is switched ON (X = 0.03), but vary the difference in the strengths of epigenetic repression between two ‘master regulators’ to draw the phase plot of population corresponding to the three states (**Fig 6C**). We noticed that for this parameter set, the population corresponding to state A changes more along the x-axis (α_AC_ - α_CA_) than along the y-axis (α_BC_ - α_CB_), i.e. the frequency of state A is more sensitive to changes in mutual epigenetic repression between A and C than those between B and C (**Fig 6C, i**). Similar trend is observed is seen for node B initially (i.e. change in the frequency of state B is more prominent along the y-axis (α_BC_ - α_CB_) than along the x-axis (α_AC_ - α_CA_)), but at higher values of α_AC_ - α_CA_, the population corresponding to state B is also affected (**Fig 6C, ii**). Because C is involved in both the instances of mutual epigenetic repression (i.e., with B and with A), the population corresponding to state C falls with an increase in incoming epigenetic repression from either node – A or B. When C is maximally repressing A and B at an epigenetic level (left bottom part of **Fig 6C, iii**), the population corresponding to state C is 80%. At the values corresponding to (maximum α_AC_, minimum α_BC_) and (minimum α_AC_, maximum α_BC_) (top left and right bottom parts of **Fig 6C,iii**), the population corresponding to state C falls to 60% and 55% respectively. At the value corresponding to (maximum α_AC_, maximum α_BC_), the population of C decreases to 15% (top right part of **Fig 56,iii**). Besides depending on the difference in corresponding α values, changes in population distribution can depend on X too (**Fig S6B-C**). These simulations performed for other parameter sets (P1-P5) reveal consistent trends (**Fig S14-S18**).

In terms of T-cell differentiation, these results imply that if the extent of epigenetic repression on one of the three ‘master regulators’ is strong enough as compared to the repression it can mediate on one or both of other two ‘master regulators’, the corresponding phenotypic frequency will decrease majorly.

## Discussion

Gaining a predictive understanding of the dynamics of cell-fate decisions is instrumental for decoding cell differentiation during development and homeostasis, and modulating it in pathological scenarios. Cell-fate decisions, including those seen in naïve helper T-cell differentiation into Th1, Th2 and Th17 cells, are the emergent outcomes of an entangled interplay of various levels of regulatory control – transcriptional (Evans and Jenner, 2013; Pillai and Jolly, 2021), translational (Liu *et al*., 2018; Sarkar *et al*., 2019), alternative splicing (Jolly *et al*., 2018; Radens *et al*., 2021), epigenetic (Wilson *et al*., 2009; Jia *et al*., 2019) and metabolic (Stark *et al*., 2019; Jia *et al*., 2021) among others. Recent efforts have begun to identify the dynamics of cellfate decisions by investigating time-course transcriptional data (Pedicini *et al*., 2010; Intosalmi *et al*., 2015; Cook and Vanderhyden, 2020; Deshmukh *et al*., 2021). However, how the different regulatory layers operating at varying time scales orchestrate coordinated cell decision-making at an individual cell and cell population level remains largely unclear.

Here, we investigate the dynamics of coupled transcriptional-epigenetic regulation in a network of three mutually repressing nodes forming a toggle triad. Analyzing the dynamics of a toggle triad can help elucidate CD4+ helper T-cell differentiation into Th1, Th2 and Th17 cells, given that each of the master regulator (T-bet, GATA3 and RORγT) can repress the other two at transcriptional and/or epigenetic levels directly or indirectly. We incorporated a simple phenomenological model to include epigenetic influence (Miyamoto *et al*., 2015) to demonstrate how varying strengths of epigenetic repression can alter the stability of the three cell states (Th1, Th2 and Th17) and consequently alter the proportion of these phenotypes in a differentiating CD4+ T-cell population. The stronger the epigenetic repression mediated by a master regulator, the higher the predominance of corresponding phenotype in a cell population.

Our model predicts that besides its strength, the duration for which epigenetic repression is ‘active’ can modulate the population heterogeneity during helper T cell differentiation. Specifically, a weaker epigenetic repression for longer times and a stronger repression for shorter times has similar outcomes; this prediction can help plan next experiments to decode T-cell differentiation as a function of varying cytokine doses and durations. Thus, unlike previous mathematical models for CD4+ T-cell differentiation mostly focused on steady-state analysis at a transcriptional level (Hong *et al*., 2011; Martinez-Sanchez *et al*., 2018; Puniya *et al*., 2018), our model incorporates the epigenetic-driven dynamics of plasticity and heterogeneity among Th1, Th2 and Th17 phenotypes in a CD4+ T-cell population.

Phenotypic heterogeneity has been reported *in silico* (Martinez-Sanchez *et al*., 2018; Puniya *et al*., 2018), *in vitro* and *in vivo* in the presence of a mixture of cytokines driving different T-cell phenotypes (Han *et al*., 2014; Becattini *et al*., 2015; DuPage and Bluestone, 2016; Eizenberg-Magar *et al*., 2017; DiToro and Basu, 2021; van Beek *et al*., 2021). Our model predicts that multistability in T-bet/GATA3/ RORγT regulatory network can allow for phenotypic switching and heterogeneity as its inherent dynamical property, similar to other regulatory networks driving ‘sibling’ cell fates (Zhou and Huang, 2011). Another dynamical feature of multistable systems is the presence of both stability (for individual phenotypes) and plasticity (among many phenotypes), and epigenetic remodeling may alter the balance between them. For instance, differences in chromatin marks may alter the propensity of an epithelial cell to switch to a mesenchymal phenotype under the influence of an inducer (Eichelberger *et al*., 2020; Jia *et al*., 2020). Similarly, a subset of Th2 cells have been shown to not express T-bet and IFNγ when stimulated under Th1 conditions (Messi *et al*., 2003). Recent “cross-polarization” experiments highlighted such “limited but detectable functional plasticity” for a population of Th1, Th2 and Th17 cells, indicating that these phenotypes represent relatively stable entities (Tortola *et al*., 2020). Epigenetic marks are considered to help maintain such stability and heritability of cell-fate decisions (Wilson *et al*., 2009), but whether epigenetic differences underlie such heterogeneity in response (stability vs. plasticity) in a cell population needs further investigation (van Beek *et al*., 2021). Preliminary evidence in Th1 cells pinpoints that permissive chromatin modifications coincide with the ability of Th1 cells to express IL-17 under Th17-polarizing conditions (Curtis *et al*., 2010), but it falls short of establishing a causative connection. Our model simulations imply that epigenetic repression driven by master regulators can influence the rate of switching from one phenotype to another, thus offering a quantitative dynamic platform to measure the stability (heritability) vs. plasticity propensities.

The balance between plasticity and stability is likely to depend on phenotype-specific global mapping of chromatin marks such as H3K4me3 and H3K27me3 that associate with activation and repression of gene expression respectively (Wei *et al*., 2009). Higher plasticity has been shown to be concurrent with the presence of bivalent chromatin (i.e. simultaneous presence of active and repressive marks) (Chaffer *et al*., 2013). For instance, the *Foxp3* promoter is not epigenetically repressed in Th17 cells, possibly enabling Th17-Treg plasticity (Wei *et al*., 2009). In our phenomenological model which does not explicitly capture the molecular details of epigenetic repression (Huang and Lei, 2019; Zhao *et al*., 2021), a bivalent chromatin state can be conceptually mapped on to the regions of relatively weak epigenetic influence of one master regulator on others. Thus, the weaker the epigenetic influence of node A on node B (α_AB_) relative to that of node B on node A (α_BA_), the higher the expected plasticity of the phenotype driven by node A. Indeed, this trend is observed in our simulations both in terms of plasticity of phenotype and the consequent population distribution. Therefore, our model can possibly explain the high plasticity observed for Th17 cells observed in many contexts such as cancer and autoimmunity (Stadhouders *et al*., 2018; Cerboni *et al*., 2021). In other words, we propose that increased plasticity of Th17 cells may be a consequence of a) weak epigenetic repression driven by RORγT on T-bet and/or GATA3, and/or b) strong epigenetic repression mediated by T-bet and/or GATA3 on RORγT.

Together, despite the limitations of investigating a minimalistic regulatory network and incorporating epigenetic influence only at a phenomenological level, our model simulations offer valuable insights into the dynamics of phenotypic plasticity and heterogeneity in a CD4+ T-cell population comprising Th1, Th2 and Th17 phenotypes. We provide a platform to quantify the plasticity and stability of different phenotypes and the overall phenotypic distribution as a function of varying strengths of epigenetic influence mediated by the master regulators (T-bet, GATA3, RORγT) on one another in a toggle triad (**Fig 6**). Various instances of plasticity among Th1, Th2 and Th17 phenotypes have been seen depending on the microenvironment (Krawczyk *et al*., 2007; Zhou *et al*., 2009; Geginat *et al*., 2016; Kanamori *et al*., 2018; Tortola *et al*., 2020), but whether this switching happens back and forth (for instance, Th1 being converted to Th2, and converting back to Th1 upon the removal of signal) remains to be investigated. The extent of such reversibility can depend on, among other factors, duration and dose of inducing signals as well as chromatin state of various regulators (Stadhouders *et al*., 2018), as seen in other cell-fate decision-making scenarios (Jia *et al*., 2019; Katsuno *et al*., 2019; Eichelberger *et al*., 2020). Our model simulations provide a framework to understand the possible conditions that may be needed for bidirectional transitions, in the form of intrinsic (epigenetic regulation) and/or extrinsic (cytokine) factors.

Our next steps include extending this T-bet/GATA3/RORγT toggle triad network to include the “master regulators” of other lineages that CD4+ T-cells can differentiate into, such as induced T-regulatory cells (iTregs) and T follicular helper (Tfh) cells among others (Martinez-Sanchez *et al*., 2018). It would be intriguing to observe what network topologies are required to explain this diversity of phenotypic repertoire of T cells. The design principles learnt through such analysis can not only reveal the dynamics of CD4+ T-cell differentiation, but also guide the design of multistable synthetic gene regulatory circuits (Santos-Moreno *et al*., 2020; Zhu *et al*., 2022).

## Author contributors

MKJ conceived and supervised research; ASD, SSM and SS performed research; all authors contributed to data analysis and interpretation, and in preparing manuscript.

## Acknowledgements

We thank Mr. Navin Vincent (IISc Bangalore) for help with the phase diagrams in Fig S1.

## Data availability

All parameter sets used here are given in Table S1. All codes used are given at the following GitHub repository: https://github.com/csbBSSE/Toggle-Triad-Epigenetics

## Materials and Methods

### RACIPE (RAndom Circuit PErturbation analysis)

RACIPE is a computational tool that investigates the emergent dynamics of a given network topology (Huang *et al*., 2017) which takes network topology as an input. Rather than specifying certain kinetic parameters of the system, RACIPE attempts to reveal all possible behaviours of the system by sampling these parameters over a range and simulating the model multiple times with varying parameter sets and initial conditions. The analysis of these results provides information on relations between the behaviour or states of the topology enabled by specific parametric spaces as well as the frequency or probability of different behaviours and states/phases of the network.

The formulation of interactions between two nodes in the network, say a node A being inhibited by a node B, in RACIPE is given by the following equation:

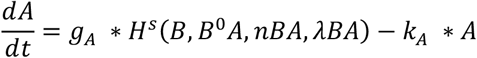

Where *g_A_* and *k_A_* are intrinsic production and degradation rates of node A respectively, and the Hill function *H^s^*(*B,B*^0^*A,nBA,λBA*) represents the interaction (here, inhibition) of node B on node A. Thus, the first term on RHS of the equation dictates the net production rate of the node A, while the second term of the equation dictates the degradation rate.

The interaction term is further expanded as,

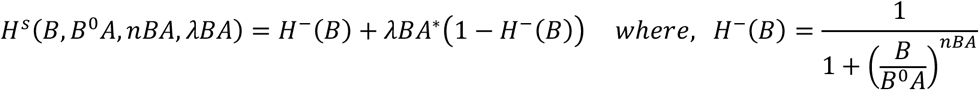

The base formulation uses hill function, but is modified to include both activation/inhibition into the same equation rather than using two separate ones. The usage of Hill functions to represent the inhibition or activation between genes is a consequence of using a biochemical rate equation formulation of gene expression (Edelstein-Keshet, 2005; Santillan, 2008). In this formulation, called as a shifted Hill function (Lu *et al*., 2013), the parameters include the threshold (*B*^0^*A*), hill coefficient (*nBA*) and fold-change value (*λBA*). The threshold determines the expression level of node B over which the inhibitory link from node B to node A is more active. Decreasing the threshold activates the link even at low expression levels of node B. The hill coefficient determines how quickly the effect of inhibition escalates with increasing expression level of node B (cooperativity) while the fold-change value determines the degree of the effect of inhibition (or activation).

Apart from the network topology as an input, the number of parameter sets and number of initial conditions per parameter set can be input; their default values are 10,000 and 100 respectively). Default values (given below) of sampling ranges for parameters can also be modified for different simulations. Default values of g and k are between (1,100) and (0.1,1) respectively; hill coefficient is sampled from the set {1,2,3,4,5,6}; Fold-change value is sampled from (1,100) for activation and (0.01,1) for inhibition. The threshold is calculated such that for all the parameter sets of the RACIPE model ensemble, each interaction has a roughly 50% chance of being functional (Huang *et al*., 2017). A parameter set is classified as enabling monostability, bistability, tristability etc. depending on the number of different steady states the 1000 initial conditions that the system converges to at the end of simulation.

For the purpose of this manuscript, we have only shortlisted parameter sets which enabled tristability with the states as {Abc, aBc, abC}. Additionally, we placed a criterion of at least 20% of the initial conditions ending up in all three of the states, to focus on parameter sets with comparable relative stability of the three states. We then selected few representative parameter sets for performing simulations shown in this manuscript.

### Mathematical framework for epigenetic feedback

The formalism used for epigenetic feedback tries to emulate the process at a phenomenological level. The referred phenomenon is that the longer a node stays at high expression, the higher the chance it has to stay high (Miyamoto *et al*., 2015), potentially because of epigenetic remodelling it is capable of ensuring which may repress its inhibitors via chromatin changes, as seen for various cell-fate decision cases (Díaz-López *et al*., 2015; Somarelli *et al*., 2016). We introduced an epigenetic parameter (α) to quantify the threshold (half-maximal) levels corresponding to influence of expression levels of one node on the other two. The higher the value of α, the lower the threshold of corresponding shifted hill function (Miyamoto *et al*., 2015). This epigenetic feedback is added to the threshold instead of hill coefficient or fold-change value because it tells us about the levels of the node which epigenetically influences its target.

The equations used are as follows:

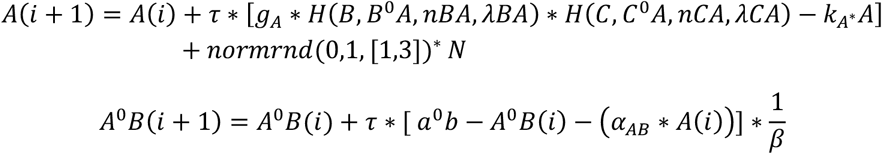

The first equation is the form of the iterative equation used for the expression level of a node. *A*(*i* + 1) and *A*(*i*) represent the node A expression levels at consecutive iterations. *τ* represents the size of time-step taken. In the equation, *g_A_* and *k_A_* are the intrinsic production and degradation rates of node A respectively. *H^s^*(*B,B*^0^*A,nBA,λBA*) and *H*(*C,C*^0^*A,nCA,λCA*) represent interaction (here, inhibition) of nodes B and C on node A respectively. The last term is the noise added to the system at set intervals in the iteration. Every time noise is added, random numbers in a vector of size 1×3 are generated from a normal distribution with mean and variance 0 and 1 respectively. (noise is added to all three nodes). These random numbers are multiplied by *N* (which is an order of magnitude lower to the mean expression level of nodes) to correspond to expression level of the nodes. Formulation of white noise term can capture different sources of biological noise such as noise due to transcriptional bursting, chromatin accessibility and protein or mRNA degradation. This formalism has previously been used to approximate biological noise (Tkacik *et al*., 2009).

The second equation represents how epigenetic feedback is employed in the formalism. Here, feedback is provided to the inhibition of node A on node B. *A*^0^*B*(*i* + 1) and *A*^0^*B*(*i*) represent threshold values corresponding to the inhibition of node B by node A at consecutive iterations. *a*^0^*b* is the set threshold value given by the chosen parameter set (i.e. without any epigenetic feedback). *α_AB_* is epigenetic parameter providing feedback corresponding to the node expression level *A*(*i*). The higher the α value, the stronger the epigenetic feedback provided or the lower is the steady state threshold value. The longer the node A is expressed high, the lower the threshold level *A*^0^*B* goes. Thus, even if the levels of node A drop due to various factors, because the threshold value is very low (representing the condition in which chromatin remodeling has taken place), the inhibition of node A on node B is still active, thus enabling node A to recover its high expression while making sure that the expression of node B remains low. *β* is a scaling factor for determine the rate of change of the threshold value and is used to control abrupt changes in node expression.

### sRACIPE

We used the webserver facility of Gene Circuit Explorer (GeneEx) to simulate stochastic dynamics of gene regulatory circuits: https://geneex.jax.org/. The tool tries to account for stochastic effects due to cell to cell variation and low copy numbers in individual cells by including a noise term based on a Wiener process (*W_t_*) with a variance. The stochastic differential equation (SDEs) are solved using Euler-Maruyama method (Kohar and Lu, 2018).

### Scoring of Th1, Th2 and Th17 gene signatures

To calculate the activity scores for specific signatures, the ssGSEA metric (Barbie *et al*., 2009) was used on gene lists of the Th1, Th2 and Th17 cell types obtained (Table S3 in (Radens *et al*., 2020)). We also computed average z-score to be used as a metric of quantification for dataset GSE62484.

