## Supplementary Figures for "Emergent dynamics of a three-node regulatory network explain phenotypic switching and heterogeneity: a case study of Th1/Th2/Th17 cell differentiation"

### Fig S1

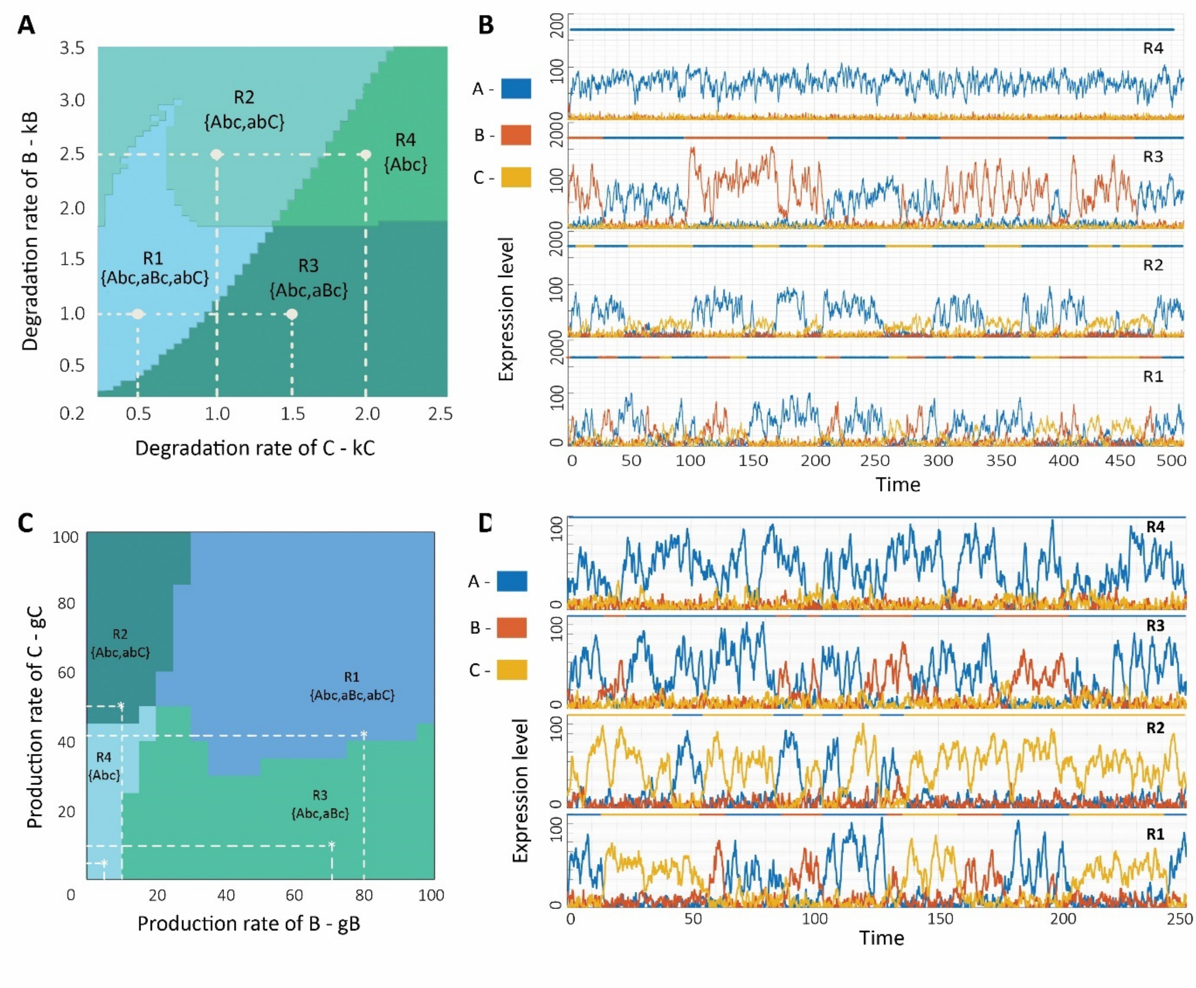


***Fig S1: A)*** *Phase diagram of toggle triad network dynamics with bifurcation parameters as degradation rates of C (kC) and B (kB). Different regions (R1: tristability – {Abc, aBc, abC}; R2: bistability – {Abc, abC}, R3: bistability – {Abc, aBC}, R4: monostability – {Abc}) are marked.* ***B)*** *Stochastic simulations of the network for various representative parameter sets (marked in B) with varying degradation rates of C and B (all other parameter values being the same) corresponding to each of the four marked regions in phase plot.* ***C)*** *Same as A) but with bifurcation parameters as production rates of C (gC) and B (gB).* ***D)*** *Same as B) but with varying production rates of C and B (all other parameter values being the same. Parameter set P1 (Table S1) is used here. Parameter values are provided in Table S1. Bars drawn above represent corresponding phenotype – Abc (state A), aBc (state B) and abC (state C).*

### Fig S2

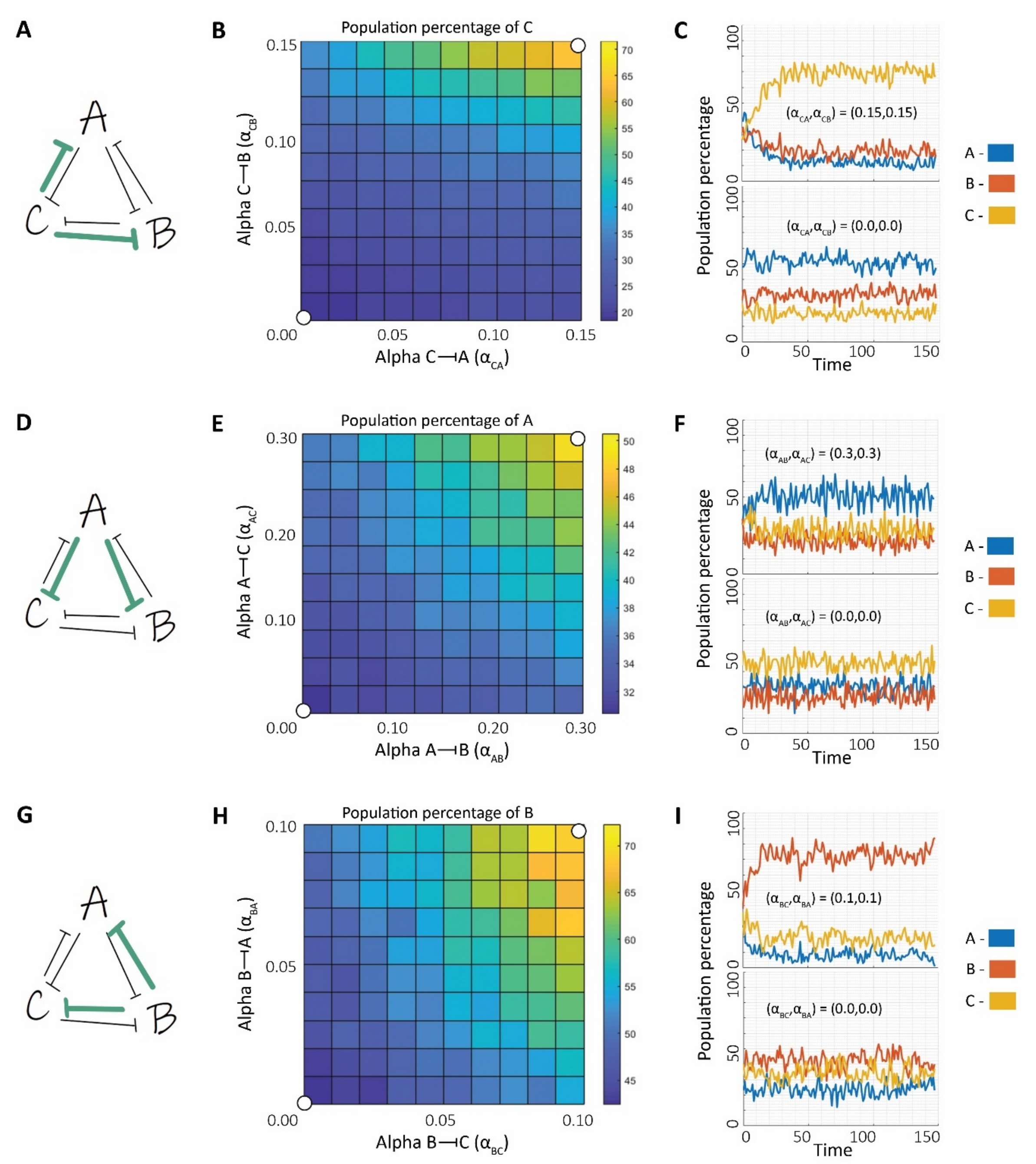


***Fig S2: A)** Toggle Triad network topology in which interactions marked in green incorporate epigenetic repression. **B)** Phase plot showing population percentage of C with bifurcation parameters as the α values corresponding to the epigenetic feedback of C -| A and C -| B. The white dots marked correspond to the dynamics simulation in **C. C)** Dynamics of distribution of population percentage between A, B and C for pairs of $\alpha_{CA}$ and $\alpha_{CB}$ values corresponding to maximum and minimum strengths of epigenetic feedback. **A, B and C** correspond to parameter set P1. **D), E)** and **F)** Same as **A, B** and **C** but for parameter set P2 and with epigenetic repression as shown in **D**. **G), H)** and **I)** Same as **A, B** and **C** but for parameter set P3 and with epigenetic repression as shown in **G**. Parameter sets given in Table S1.*

### Fig S3

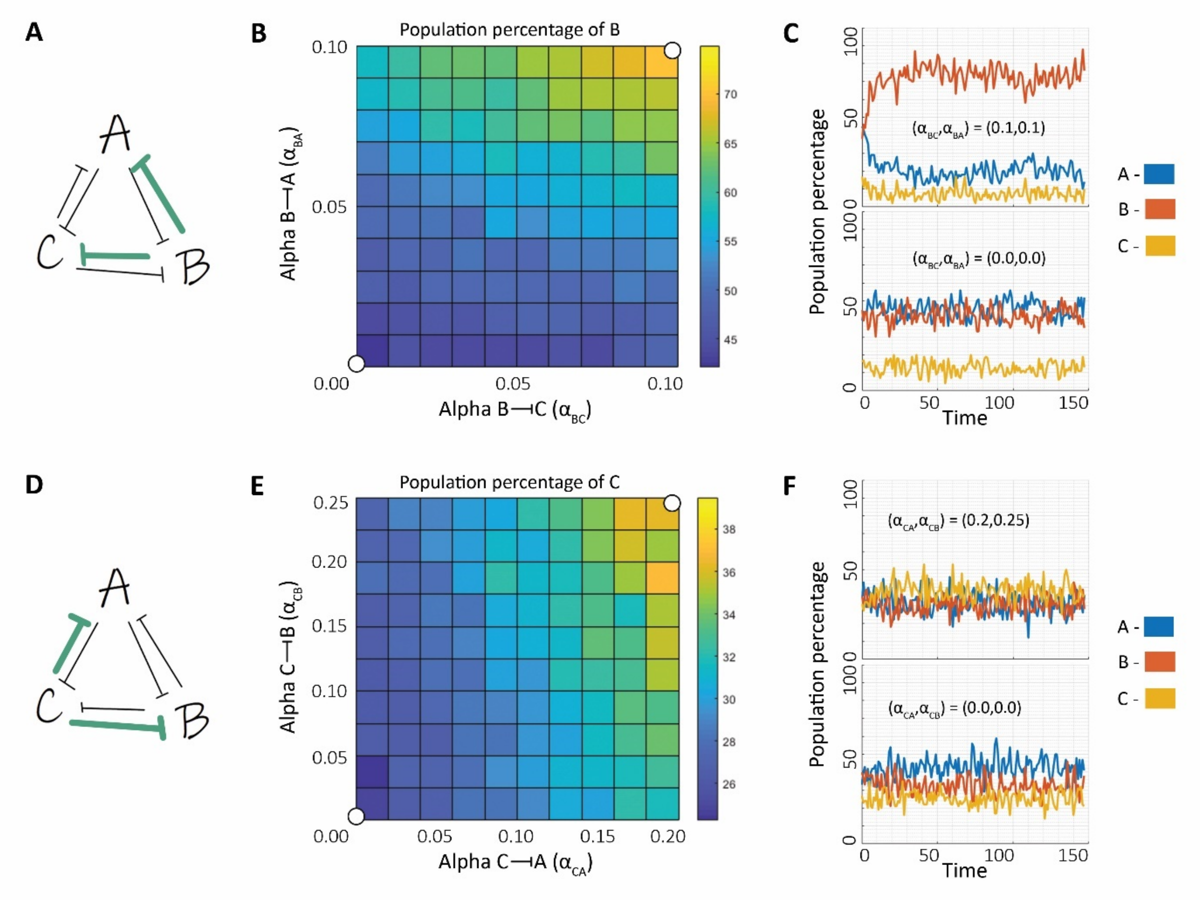


***Fig S3: A)*** *Toggle Triad network topology in which interactions marked in green are being provided with epigenetic feedback* ***B)*** *Phase plot showing population percentage of B with bifurcation parameters as the α values corresponding to epigenetic repression of B -| A and B -| C. White dots marked correspond to the dynamics simulation in* ***C. C)*** *Dynamics of distribution of population percentage between A, B and C for pairs of $\alpha_{BA}$ and $\alpha_{BC}$ values corresponding to maximum and minimum strengths of epigenetic repression.* ***A, B and C*** *correspond to parameter set P4.* ***D), E)*** *and* ***F)*** *Same as* ***A, B*** *and* ***C*** *but for parameter set P5 and with epigenetic repression as shown in* ***D****. Parameter sets given in Table S1.*

### Fig S4

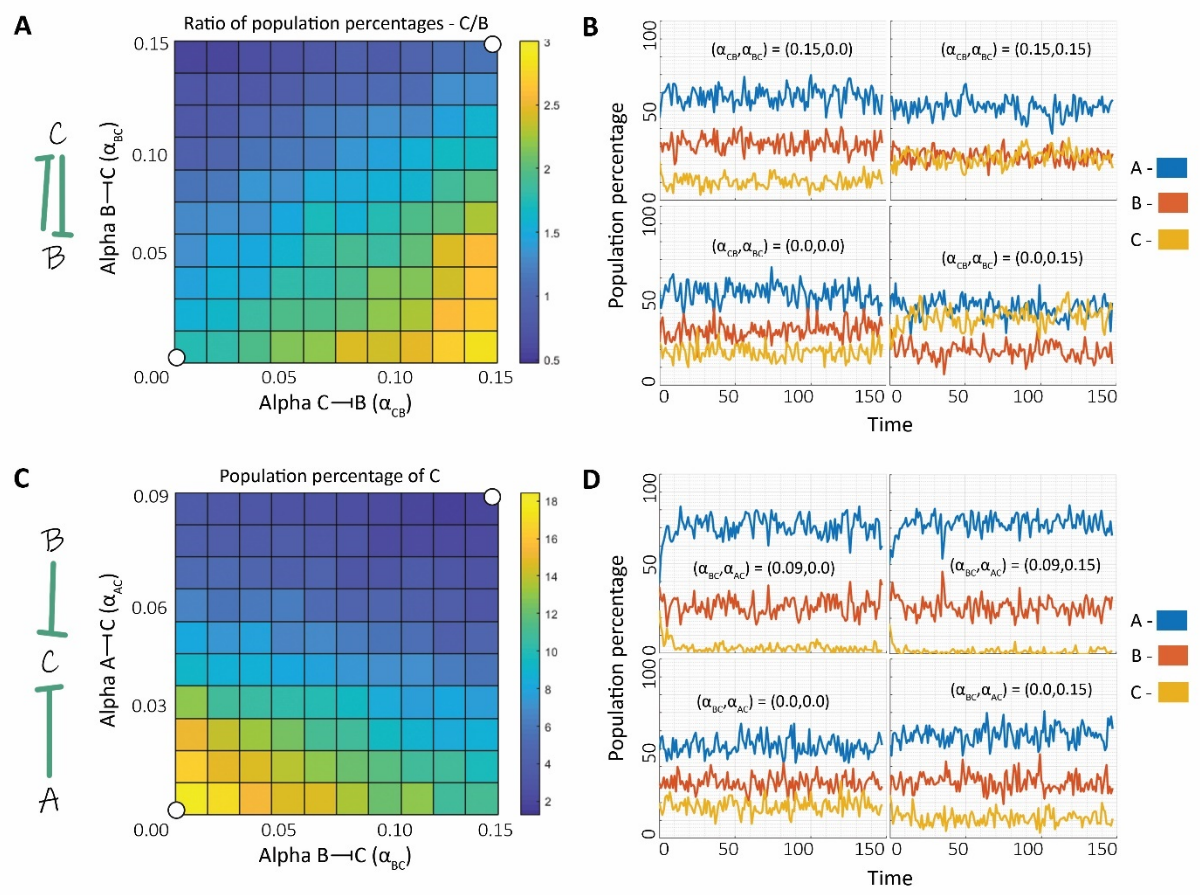


***Fig S4: A)*** *Toggle Triad network topology in which interactions marked in green are being provided with epigenetic feedback as well as phase plot showing ratio of population percentage of C to that of B with bifurcation parameters as the α values corresponding to the epigenetic feedback of C-|B and B-|C. The white dots marked correspond to the dynamics simulation in* ***B. B)*** *Dynamics of distribution of population percentage between A, B and C for certain pairs of* $\alpha_{BC}$ *and* $\alpha_{CB}$ *values.* ***C)*** *Toggle Triad network topology in which interactions marked in green are being provided with epigenetic feedback as well as phase plot showing population percentage of C with bifurcation parameters as the α values corresponding to the epigenetic feedback of A-|C and B-|C. The white dots marked correspond to the dynamics simulation in* ***D. D)*** *Dynamics of distribution of population percentage between A, B and C for certain pairs of* $\alpha_{AC}$ *and* $\alpha_{BC}$ *values. Parameter set P1 (Table S1) is used here.*

### Fig S5

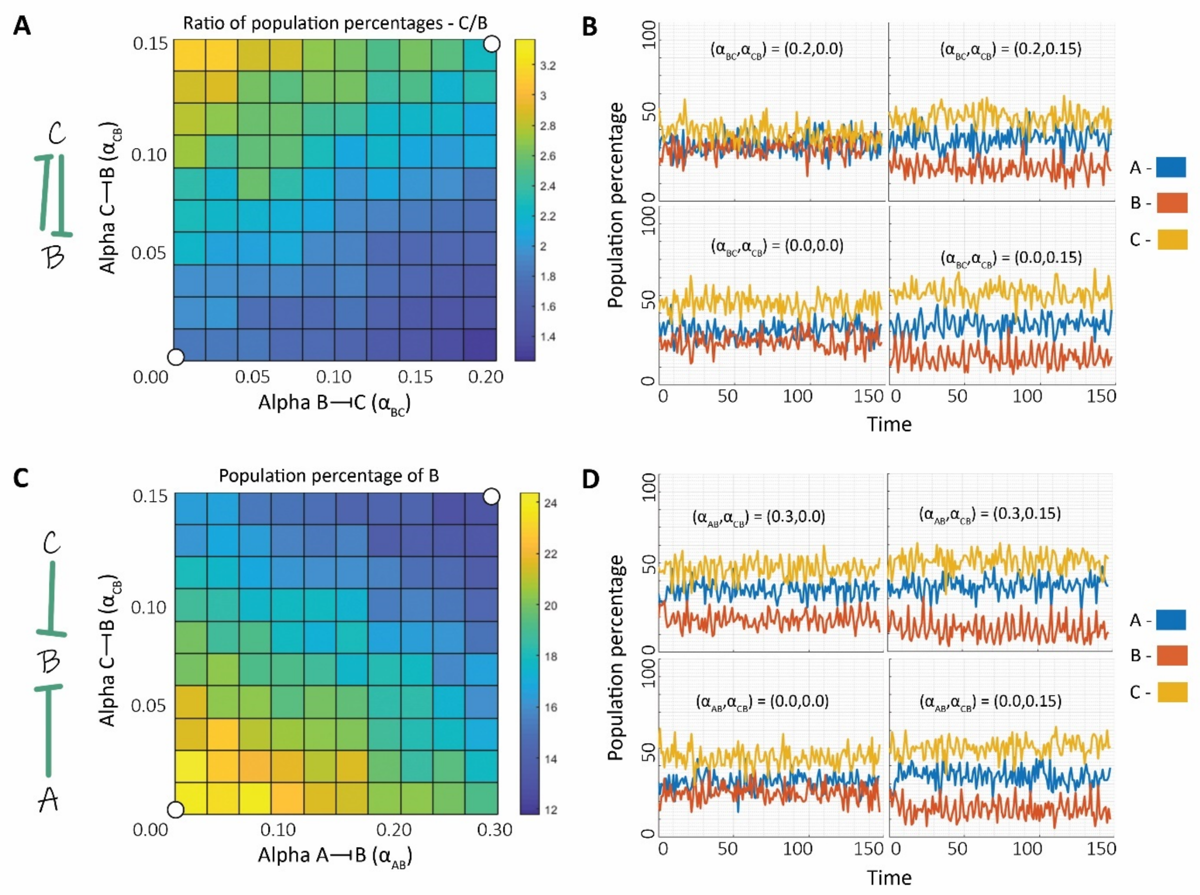


***Fig S5: A)*** *Toggle Triad network topology in which interactions marked in green are being provided with epigenetic feedback as well as phase plot showing ratio of population percentage of C to that of B with bifurcation parameters as the α values corresponding to the epigenetic feedback of C-|B and B-|C. The white dots marked correspond to the dynamics simulation in* ***B. B)*** *Dynamics of distribution of population percentage between A, B and C for certain pairs of* $\alpha_{BC}$ *and* $\alpha_{CB}$ *values.* ***C)*** *Toggle Triad network topology in which interactions marked in green are being provided with epigenetic feedback as well as phase plot showing population percentage of B with bifurcation parameters as the α values corresponding to the epigenetic feedback of A-|B and C-|B. The white dots marked correspond to the dynamics simulation in* ***D. D)*** *Dynamics of distribution of population percentage between A, B and C for certain pairs of* $\alpha_{AB}$ *and* $\alpha_{CB}$ *values. Parameter set P2 (Table S1) is used here.*

### Fig S6

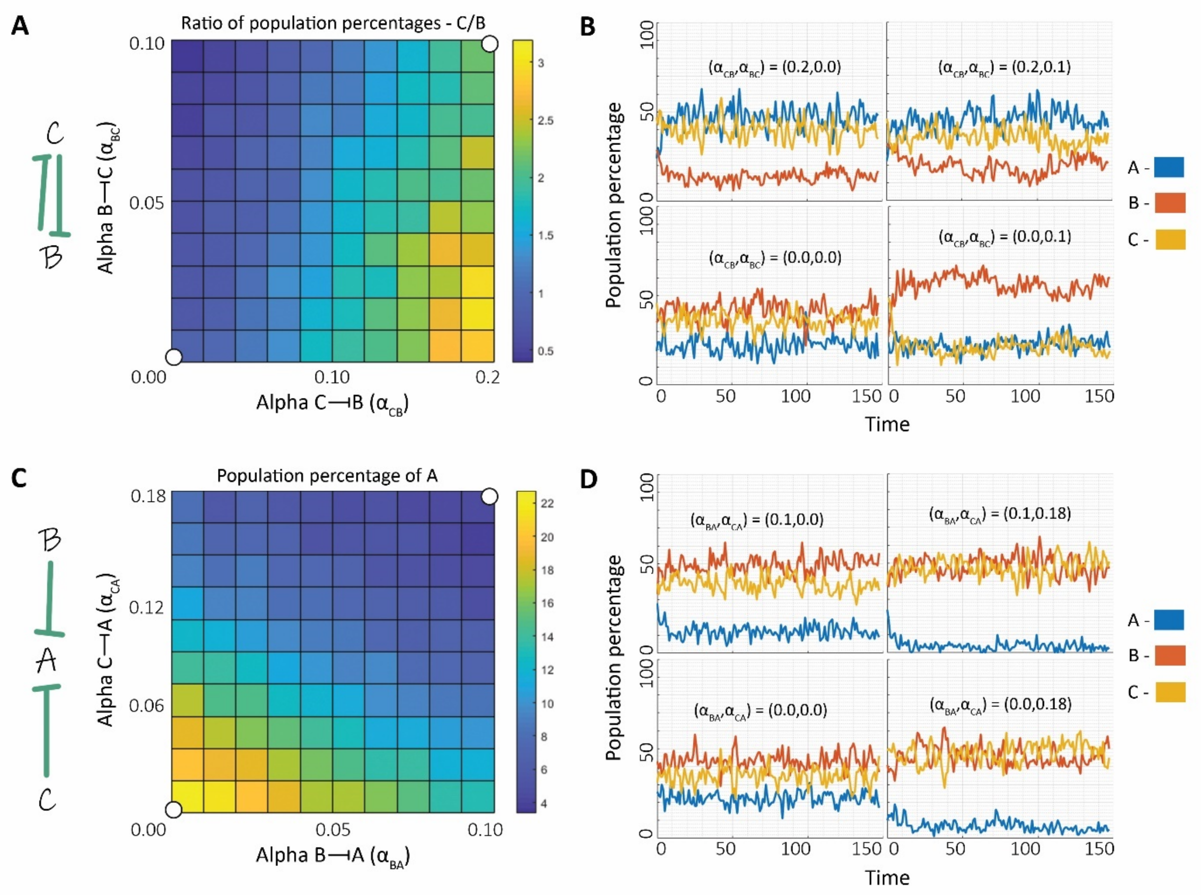


***Fig S6: A)*** *Toggle Triad network topology in which interactions marked in green are being provided with epigenetic feedback as well as phase plot showing ratio of population percentage of C to that of B with bifurcation parameters as the α values corresponding to the epigenetic feedback of C-|B and B-|C. The white dots marked correspond to the dynamics simulation in* ***B. B)*** *Dynamics of distribution of population percentage between A, B and C for certain pairs of $\alpha_{BC}$ and $\alpha_{CB}$ values.* ***C)*** *Toggle Triad network topology in which interactions marked in green are being provided with epigenetic feedback as well as phase plot showing population percentage of A with bifurcation parameters as the α values corresponding to the epigenetic feedback of B-|A and C-|A. The white dots marked correspond to the dynamics simulation in* ***D. D)*** *Dynamics of distribution of population percentage between A, B and C for certain pairs of αBA and αCA values. Parameter set P3 (Table S1) is used here.*

### Fig S7

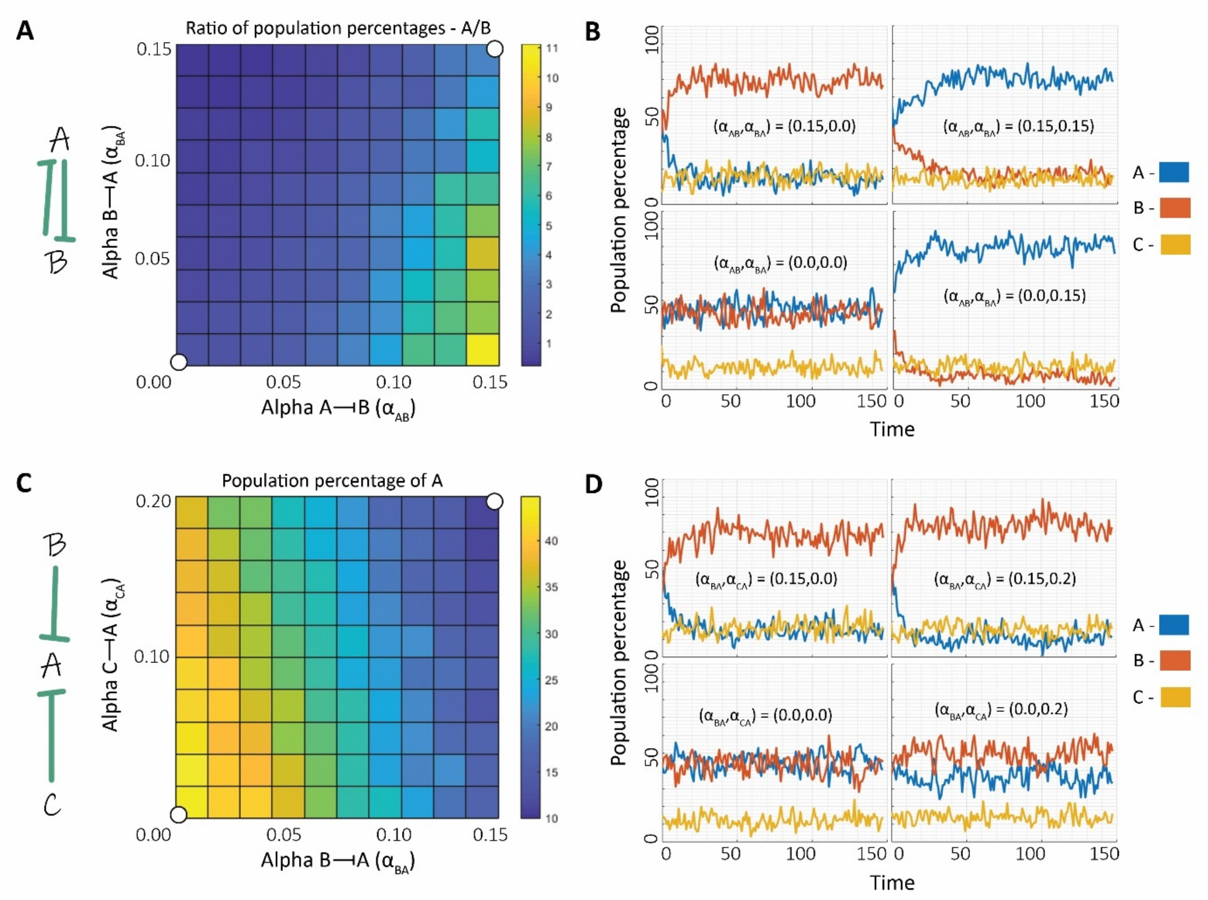


***Fig S7: A)*** *Toggle Triad network topology in which interactions marked in green are being provided with epigenetic feedback as well as phase plot showing ratio of population percentage of A to that of B with bifurcation parameters as the α values corresponding to the epigenetic feedback of A-|B and B-|A. The white dots marked correspond to the dynamics simulation in* ***B. B)*** *Dynamics of distribution of population percentage between A, B and C for certain pairs of* $\alpha_{AB}$ *and* $\alpha_{BA}$ *values.* ***C)*** *Toggle Triad network topology in which interactions marked in green are being provided with epigenetic feedback as well as phase plot showing population percentage of A with bifurcation parameters as the α values corresponding to the epigenetic feedback of B-|A and C-|A. The white dots marked correspond to the dynamics simulation in* ***D. D)*** *Dynamics of distribution of population percentage between A, B and C for certain pairs of αBA and αCA values. Parameter set P4 (Table S1) is used here.*

### Fig S8

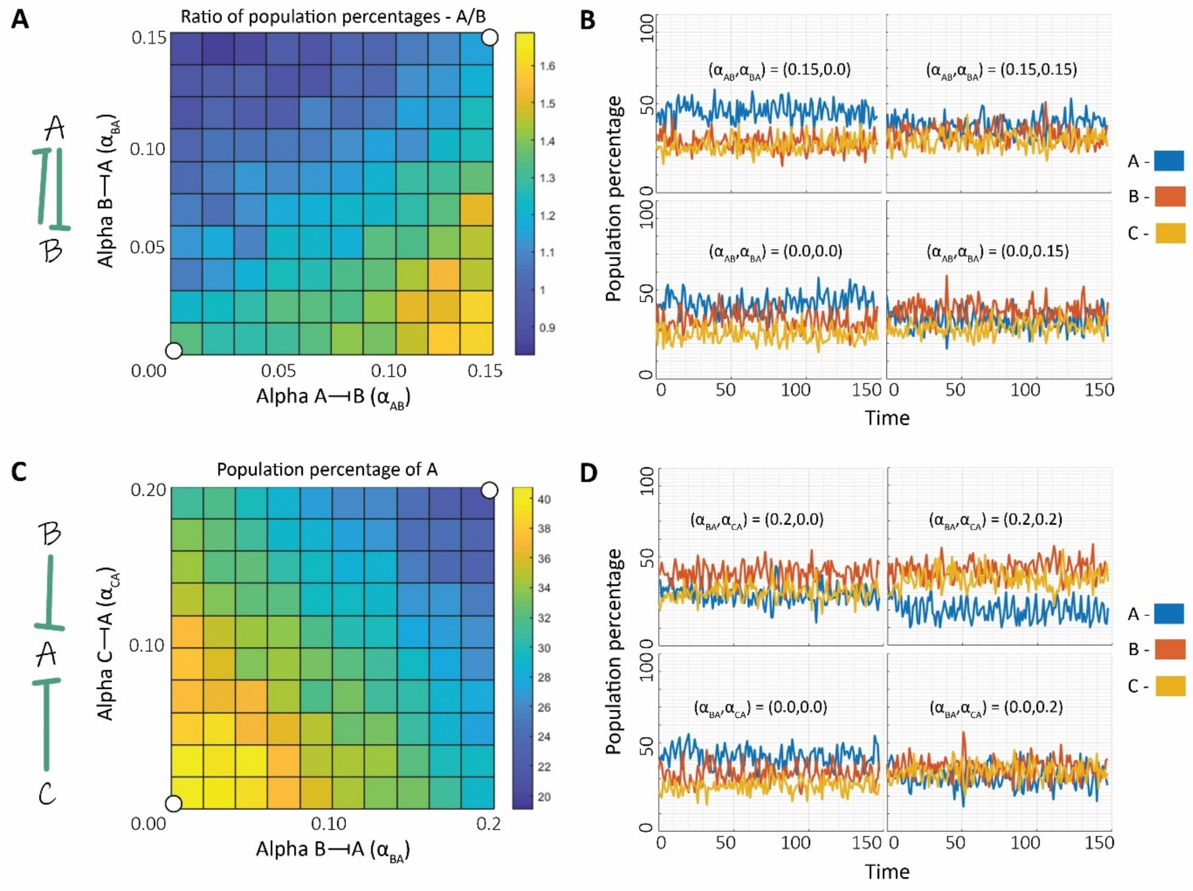


***Fig S8: A)*** *Toggle Triad network topology in which interactions marked in green are being provided with epigenetic feedback as well as phase plot showing ratio of population percentage of A to that of B with bifurcation parameters as the α values corresponding to the epigenetic feedback of A-|B and B-|A. The white dots marked correspond to the dynamics simulation in* ***B. B)*** *Dynamics of distribution of population percentage between A, B and C for certain pairs of* $\alpha_{AB}$ *and* $\alpha_{BA}$ *values.* ***C)*** *Toggle Triad network topology in which interactions marked in green are being provided with epigenetic feedback as well as phase plot showing population percentage of A with bifurcation parameters as the α values corresponding to the epigenetic feedback of B-|A and C-|A. The white dots marked correspond to the dynamics simulation in* ***D. D)*** *Dynamics of distribution of population percentage between A, B and C for certain pairs of αBA and αCA values. Parameter set P5 (Table S1) is used here.*

### Fig S9

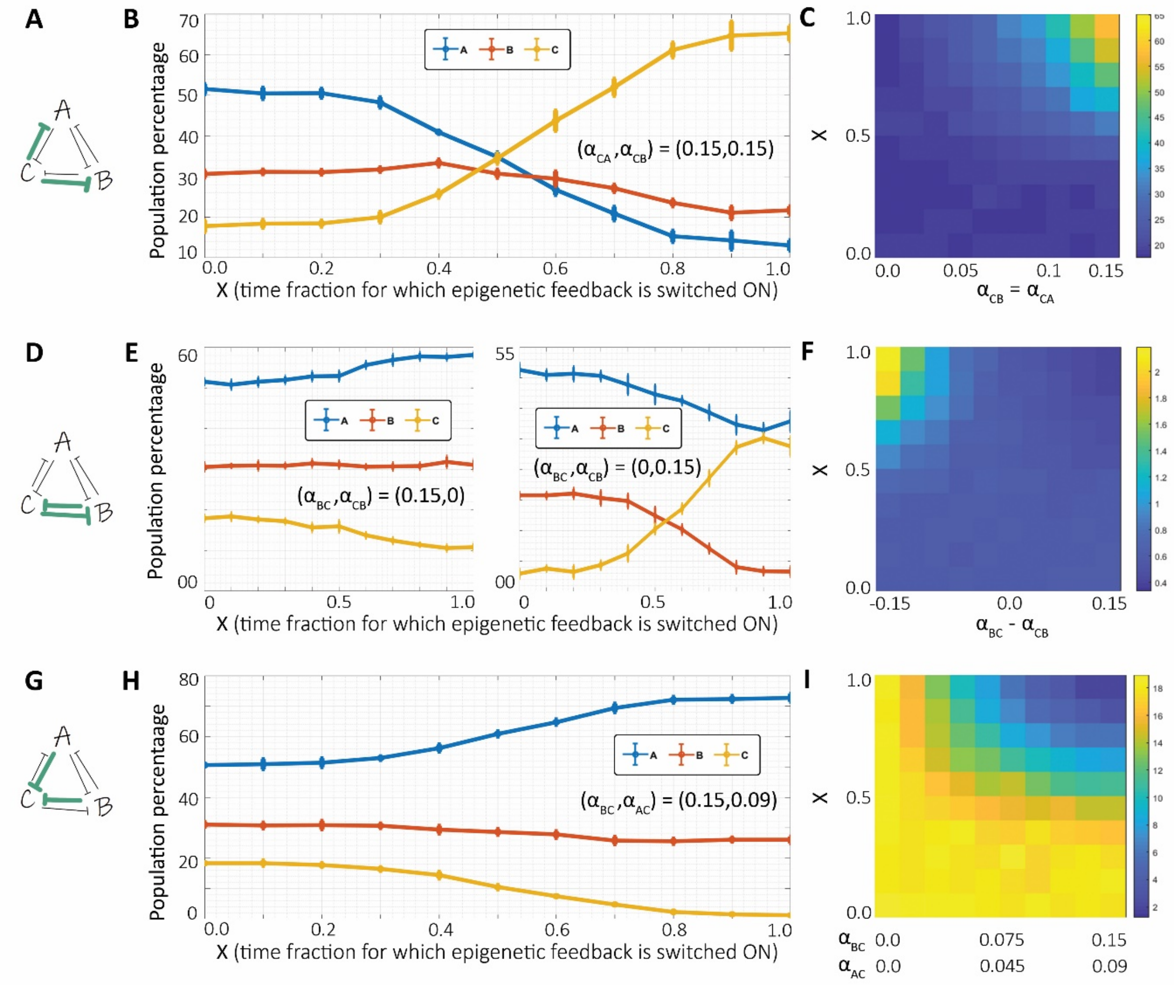


***Fig S9: A)*** *Toggle Triad network topology in which interactions marked in green are being provided with epigenetic feedback.* ***B)*** *Population percentages of A, B and C as X, the fraction of time for which the epigenetic feedback for both marked interactions is switched ON and then turned OFF.* ***C)*** *Phase plot showing population percentage of C (node from which interactions with epigenetic feedback originate) with bifurcation parameters as the α value corresponding to the epigenetic feedback of C-|A and C-|B (both same so considered on single axis) and X.* ***D)*** *Same as A)* ***E)*** *Same as B) but for two cases where feedback for one of the interactions, B-|C (C-|B) is switched ON with the other C-|B (B-|C) switched OFF.* ***F)*** *Phase plot showing ratio of population percentage of C to B (nodes between which interactions with epigenetic feedback are present) with bifurcation parameters as the difference of α values corresponding to the epigenetic feedback of B-|C and C-|B and X.* ***G)*** *Same as A).* ***H)*** *Same as B)* ***I)*** *Phase plot showing population percentage of C (node onto which interactions with epigenetic feedback terminate) with bifurcation parameters as the α value corresponding to the epigenetic feedback of A-|C and B-|C (both same so considered on single axis) and X. Parameter set P1 (Table S1) has been used here.*

### Fig S10

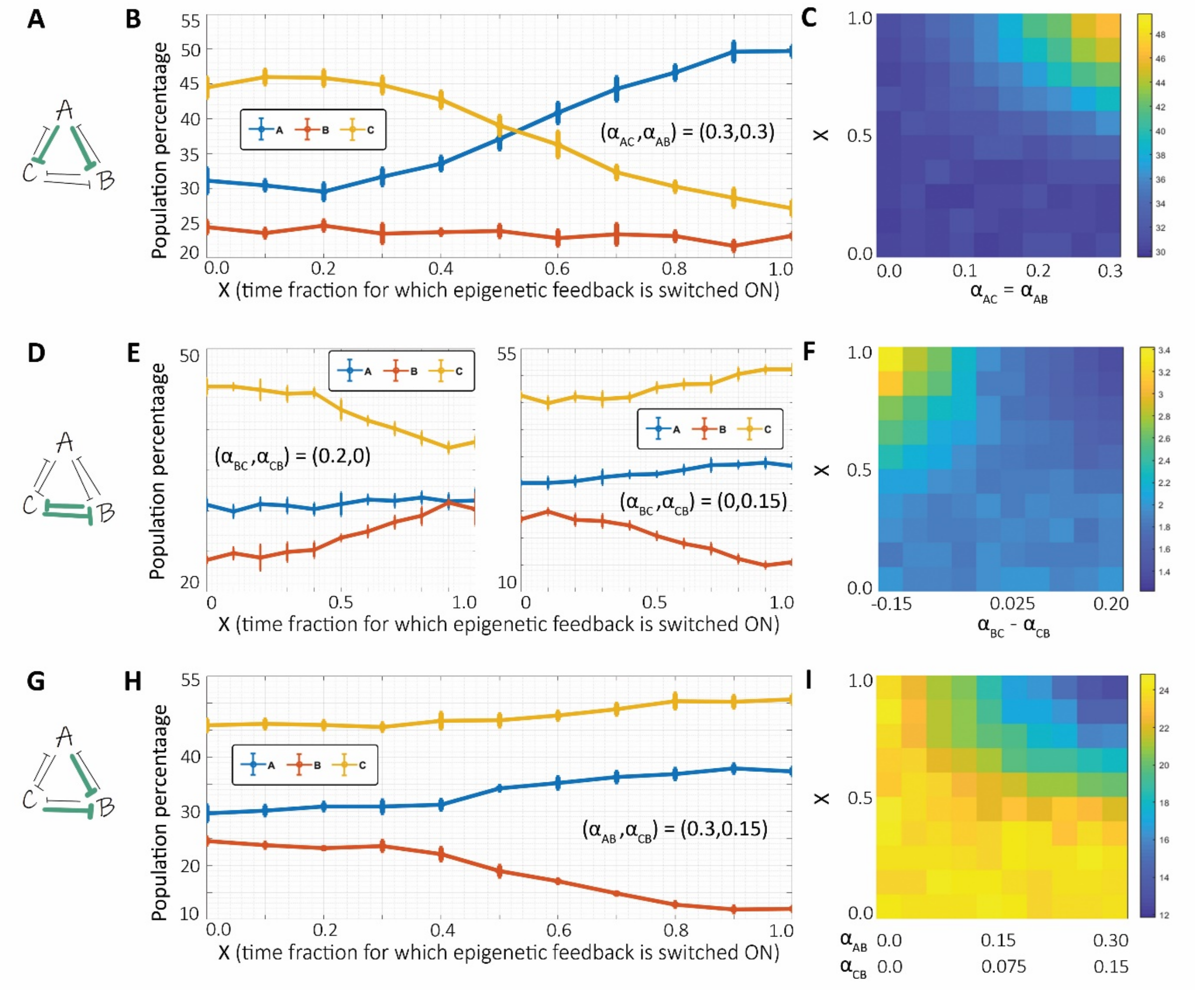


***Fig S10: A)*** *Toggle Triad network topology in which interactions marked in green are being provided with epigenetic feedback.* ***B)*** *Population percentages of A, B and C as X, the fraction of time for which the epigenetic feedback for both marked interactions is switched ON and then turned OFF.* ***C)*** *Phase plot showing population percentage of A (node from which interactions with epigenetic feedback originate) with bifurcation parameters as the α value corresponding to the epigenetic feedback of A-|B and A-|C (both same so considered on single axis) and X.* ***D)*** *Same as A)* ***E)*** *Same as B) but for two cases where feedback for one of the interactions, B-|C (C-|B) is switched ON with the other C-|B (B-|C) switched OFF.* ***F)*** *Phase plot showing ratio of population percentage of C to B (nodes between which interactions with epigenetic feedback are present) with bifurcation parameters as the difference of α values corresponding to the epigenetic feedback of B-|C and C-|B and X.* ***G)*** *Same as A).* ***H)*** *Same as B)* ***I)*** *Phase plot showing population percentage of B (node onto which interactions with epigenetic feedback terminate) with bifurcation parameters as the α value corresponding to the epigenetic feedback of A-|B and C-|B (both same so considered on single axis) and X. Parameter set P2 (Table S1) has been used here.*

### Fig S11

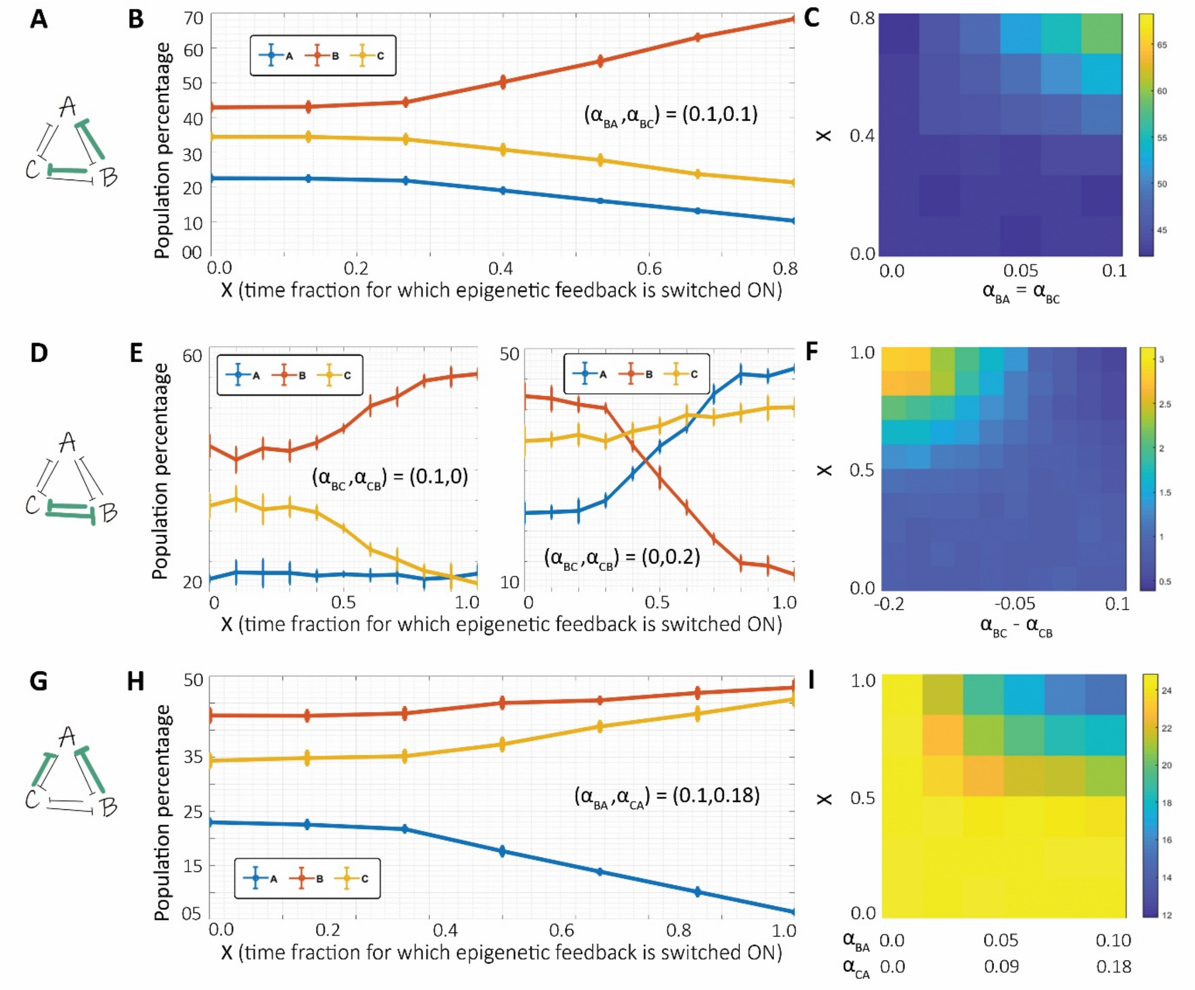


***Fig S11: A)*** *Toggle Triad network topology in which interactions marked in green are being provided with epigenetic feedback.* ***B)*** *Population percentages of A, B and C as X, the fraction of time for which the epigenetic feedback for both marked interactions is switched ON and then turned OFF.* ***C)*** *Phase plot showing population percentage of B (node from which interactions with epigenetic feedback originate) with bifurcation parameters as the α value corresponding to the epigenetic feedback of B-|A and B-|C (both same so considered on single axis) and X.* ***D)*** *Same as A)* ***E)*** *Same as B) but for two cases where feedback for one of the interactions, B-|C (C-|B) is switched ON with the other C-|B (B-|C) switched OFF.* ***F)*** *Phase plot showing ratio of population percentage of C to B (nodes between which interactions with epigenetic feedback are present) with bifurcation parameters as the difference of α values corresponding to the epigenetic feedback of B-|C and C-|B and X.* ***G)*** *Same as A).* ***H)*** *Same as B)* ***I)*** *Phase plot showing population percentage of A (node onto which interactions with epigenetic feedback terminate) with bifurcation parameters as the α value corresponding to the epigenetic feedback of B-|A and C-|A (both same so considered on single axis) and X. Parameter set P3 (Table S1) has been used here.*

Fig S12

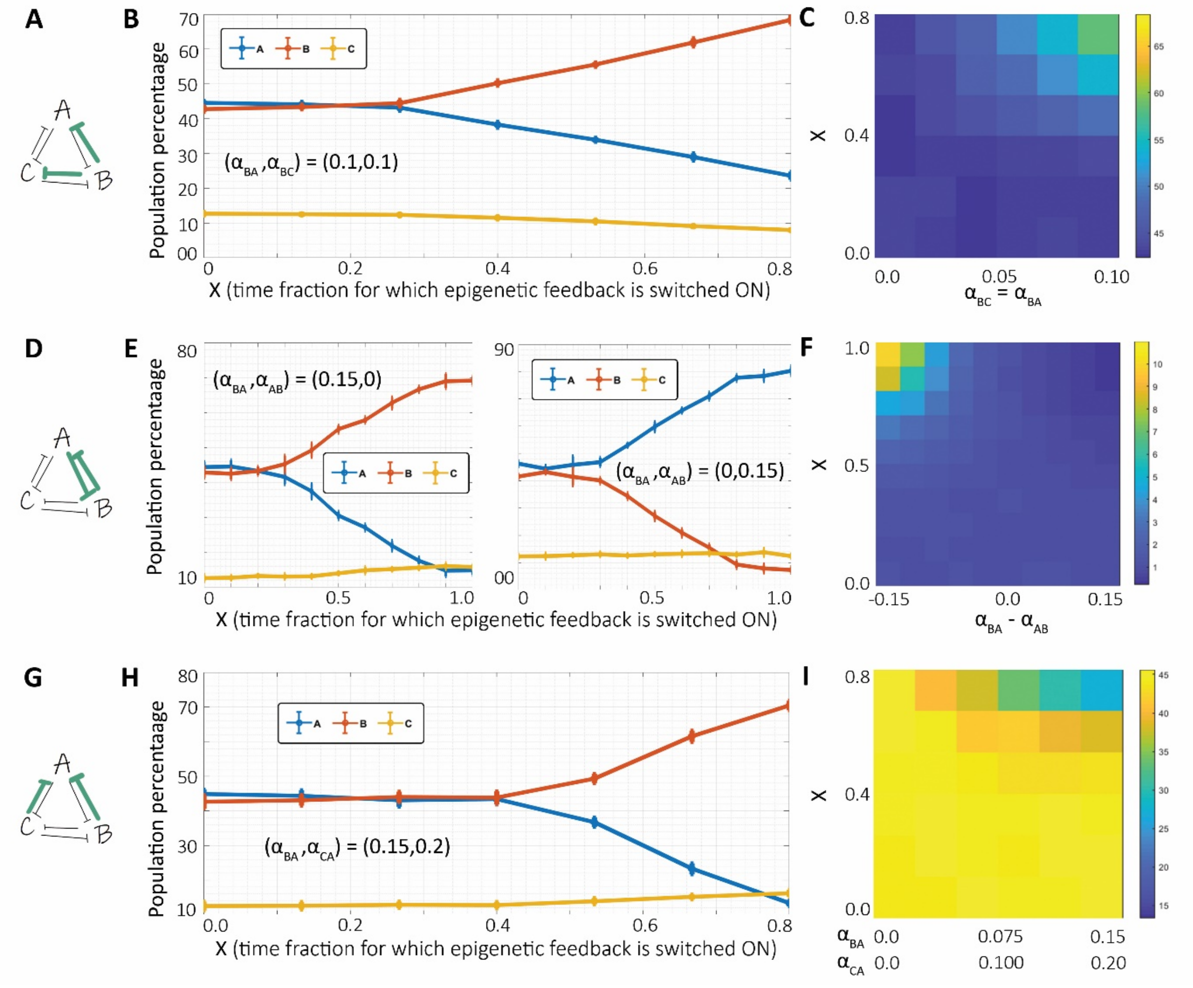


***Fig S12: A)** Toggle Triad network topology in which interactions marked in green are being provided with epigenetic feedback. **B)** Population percentages of A, B and C as X, the fraction of time for which the epigenetic feedback for both marked interactions is switched ON and then turned OFF. **C)** Phase plot showing population percentage of B (node from which interactions with epigenetic feedback originate) with bifurcation parameters as the α value corresponding to the epigenetic feedback of B-|A and B-|C (both same so considered on single axis) and X. **D)** Same as A) **E)** Same as B) but for two cases where feedback for one of the interactions, A-|B (B-|A) is switched ON with the other B-|A (A-|B) switched OFF. **F)** Phase plot showing ratio of population percentage of A to B (nodes between which interactions with epigenetic feedback are present) with bifurcation parameters as the difference of α values corresponding to the epigenetic feedback of A-|B and B-|A and X. **G)** Same as A). **H)** Same as B) **I)** Phase plot showing population percentage of A (node onto which interactions with epigenetic feedback terminate) with bifurcation parameters as the α value corresponding to the epigenetic feedback of B-|A and C-|A (both same so considered on single axis) and X. Parameter set P4 (Table S1) has been used here.*

### Fig S13

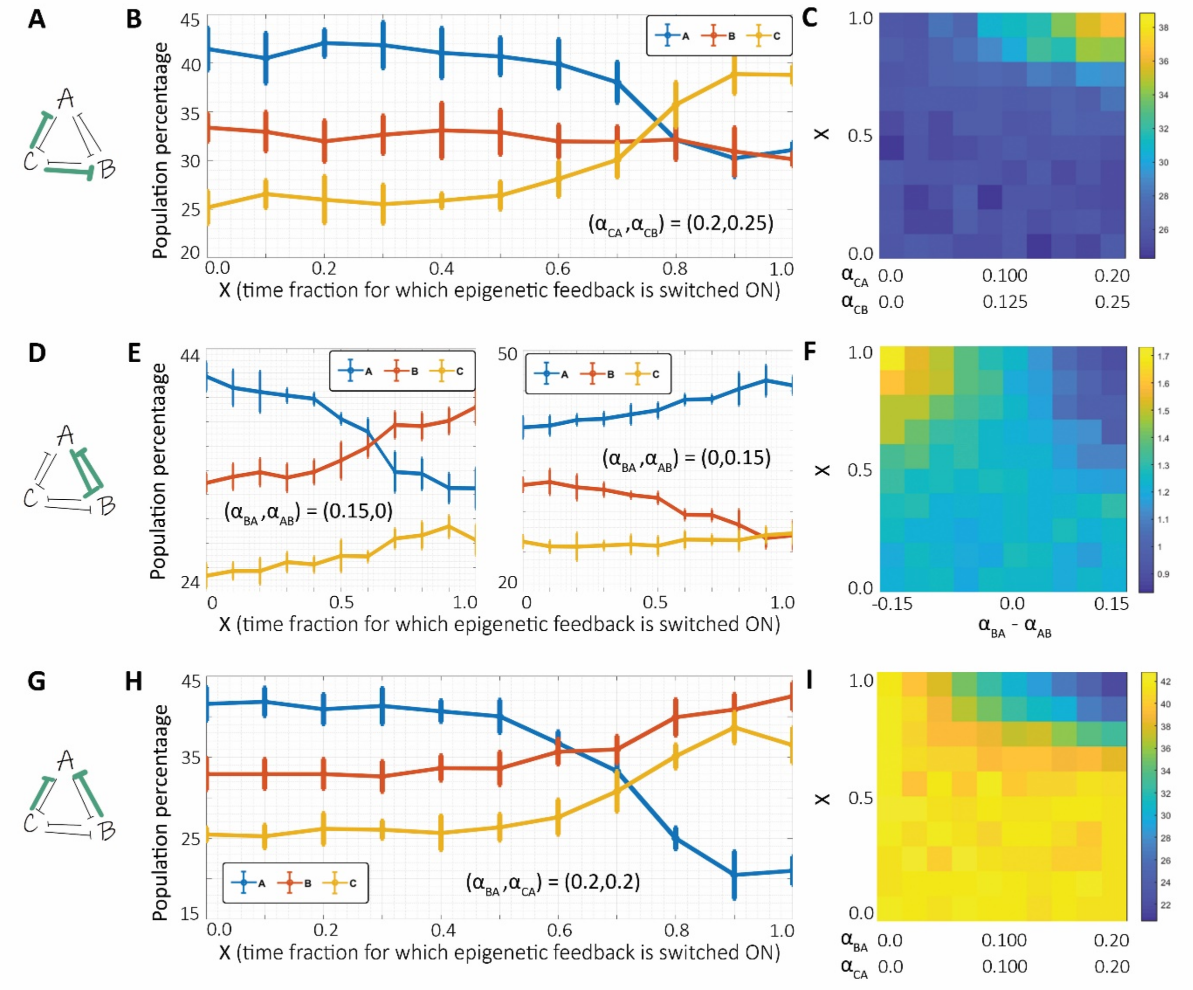


***Fig S13: A)*** *Toggle Triad network topology in which interactions marked in green are being provided with epigenetic feedback.* ***B)*** *Population percentages of A, B and C as X, the fraction of time for which the epigenetic feedback for both marked interactions is switched ON and then turned OFF.* ***C)*** *Phase plot showing population percentage of C (node from which interactions with epigenetic feedback originate) with bifurcation parameters as the α value corresponding to the epigenetic feedback of C-|A and C-|B (both same so considered on single axis) and X.* ***D)*** *Same as A)* ***E)*** *Same as B) but for two cases where feedback for one of the interactions, A-|B (B-|A) is switched ON with the other B-|A (A-|B) switched OFF.* ***F)*** *Phase plot showing ratio of population percentage of A to B (nodes between which interactions with epigenetic feedback are present) with bifurcation parameters as the difference of α values corresponding to the epigenetic feedback of A-|B and B-|A and X.* ***G)*** *Same as A).* ***H)*** *Same as B)* ***I)*** *Phase plot showing population percentage of A (node onto which interactions with epigenetic feedback terminate) with bifurcation parameters as the α value corresponding to the epigenetic feedback of B-|A and C-|A (both same so considered on single axis) and X. Parameter set P5 (Table S1) has been used here.*

### Fig S14

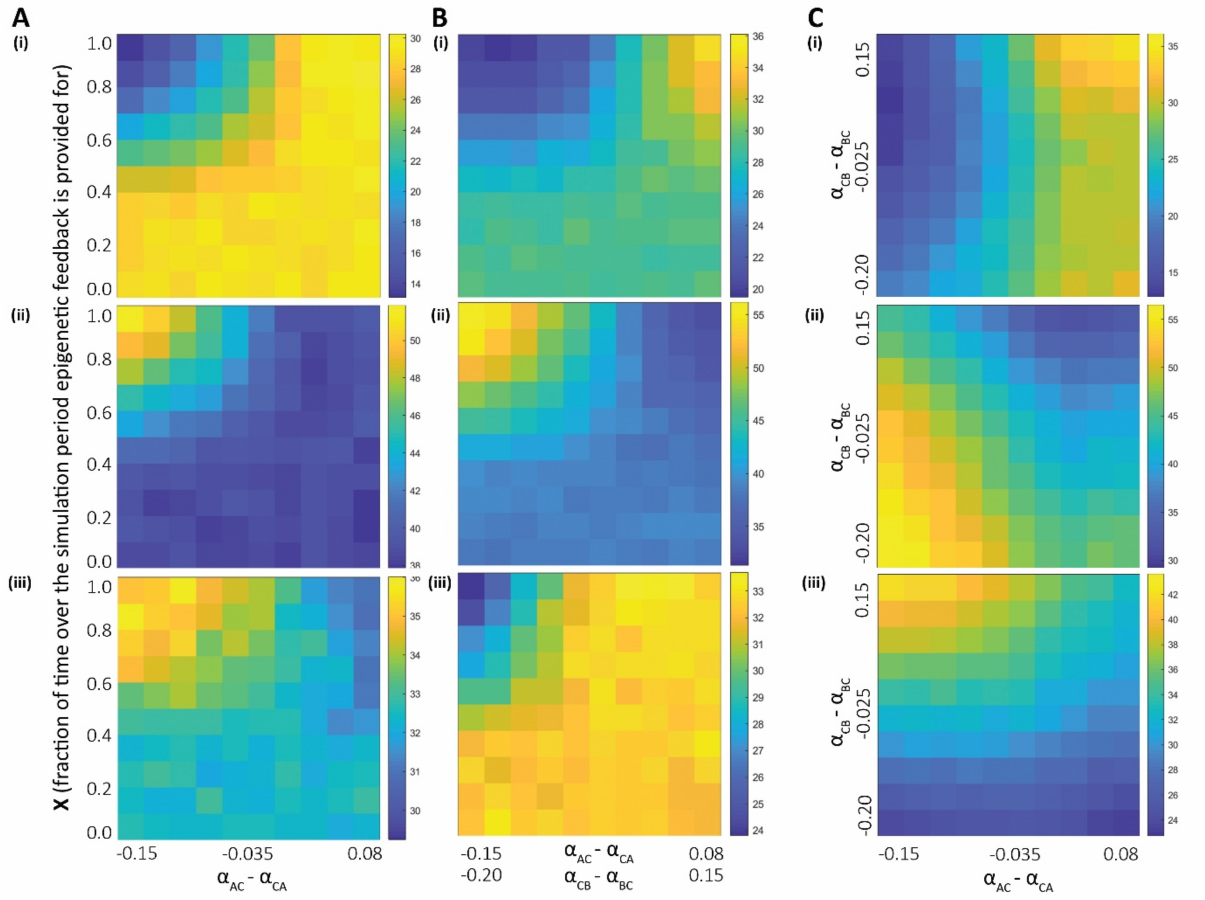


***Fig S14: A) i)*** *Phase plot of population percentage corresponding to node A with variations in two parameters: X and the relative epigenetic influence which is varied from a case of stronger repression from C to A ($\alpha_{AC} < \alpha_{CA}$) to a stronger repression from A to C ($\alpha_{AC} > \alpha_{CA}$)* ***ii)*** *Same as* ***i)*** *but for state B.* ***iii)*** *Same as* ***i)*** *but for state C.* ***B) i)*** *Same as A;* ***i)*** *but for varying relative epigenetic influence in both feedback loops (between A and C, and between B and C).* ***ii)*** *Same as* ***i)*** *but for state B.* ***iii)*** *Same as* ***i)*** *but for state C.* ***C) i)*** *Phase plot of population percentage corresponding to state A with bifurcation parameters as the difference in α values corresponding to mutual epigenetic repression ($\alpha_{CB}$ - $\alpha_{BC}$ and $\alpha_{AC}$ - $\alpha_{CA}$), at X = 1.* ***ii)*** *Same as* ***i)*** *but for state B.* ***iii)*** *Same as* ***i)*** *but for state C. Parameter set P1 (Table S1) is used here.*

### Fig S15

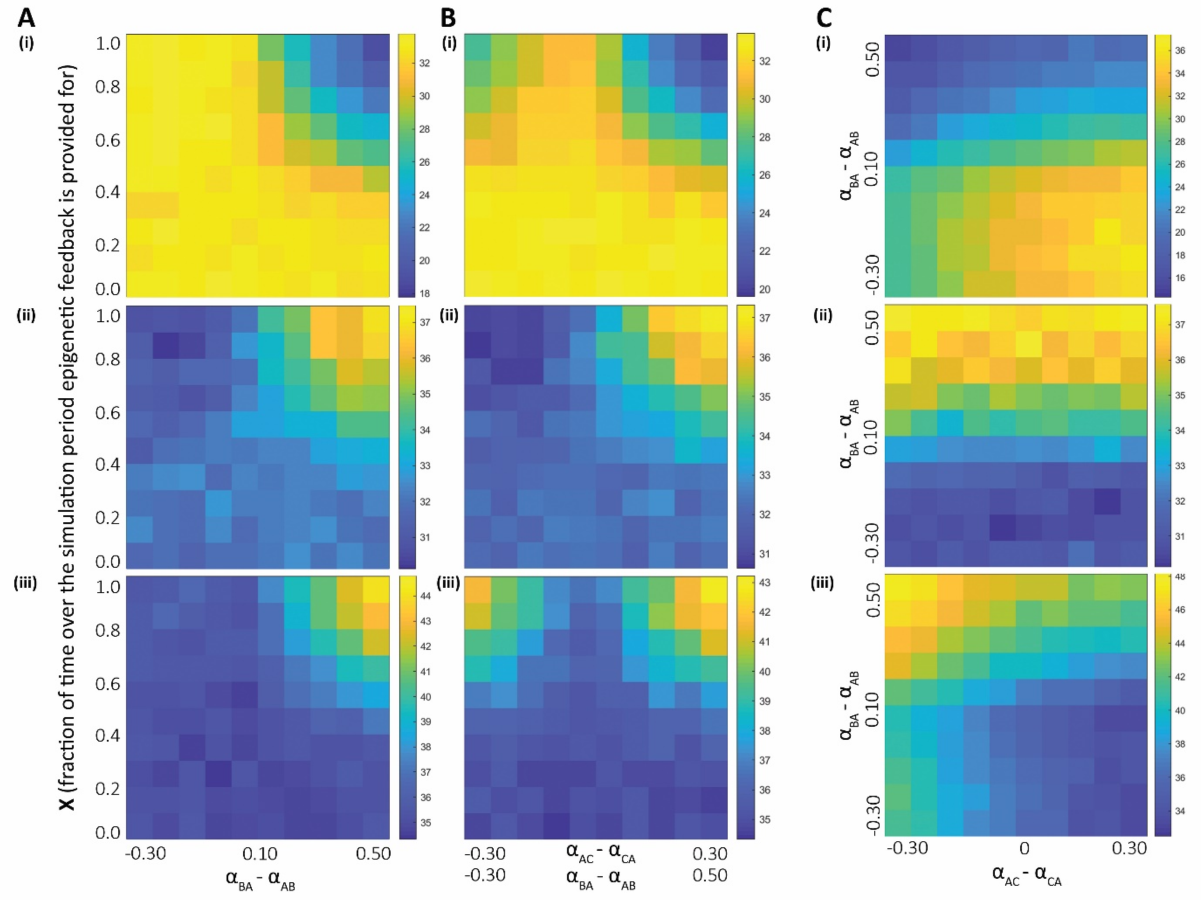


***Fig S15: A) i)*** *Phase plot of population percentage corresponding to node A with variations in two parameters: X and the relative epigenetic influence which is varied from a case of stronger repression from A to B ($\alpha_{BA} < \alpha_{AB}$) to a stronger repression from B to A ($\alpha_{BA} > \alpha_{AB}$)* ***ii)*** *Same as* ***i)*** *but for state B.* ***iii)*** *Same as* ***i)*** *but for state C.* ***B) i)*** *Same as A;* ***i)*** *but for varying relative epigenetic influence in both feedback loops (between B and A, and between C and A).* ***ii)*** *Same as* ***i)*** *but for state B.* ***iii)*** *Same as* ***i)*** *but for state C.* ***C) i)*** *Phase plot of population percentage corresponding to state A with bifurcation parameters as the difference in α values corresponding to mutual epigenetic repression ($\alpha_{BA} - \alpha_{AB}$ and $\alpha_{AC} - \alpha_{CA}$), at X = 1.* ***ii)*** *Same as* ***i)*** *but for state B.* ***iii)*** *Same as* ***i)*** *but for state C. Parameter set P2 (Table S1) is used here.*

### Fig S16

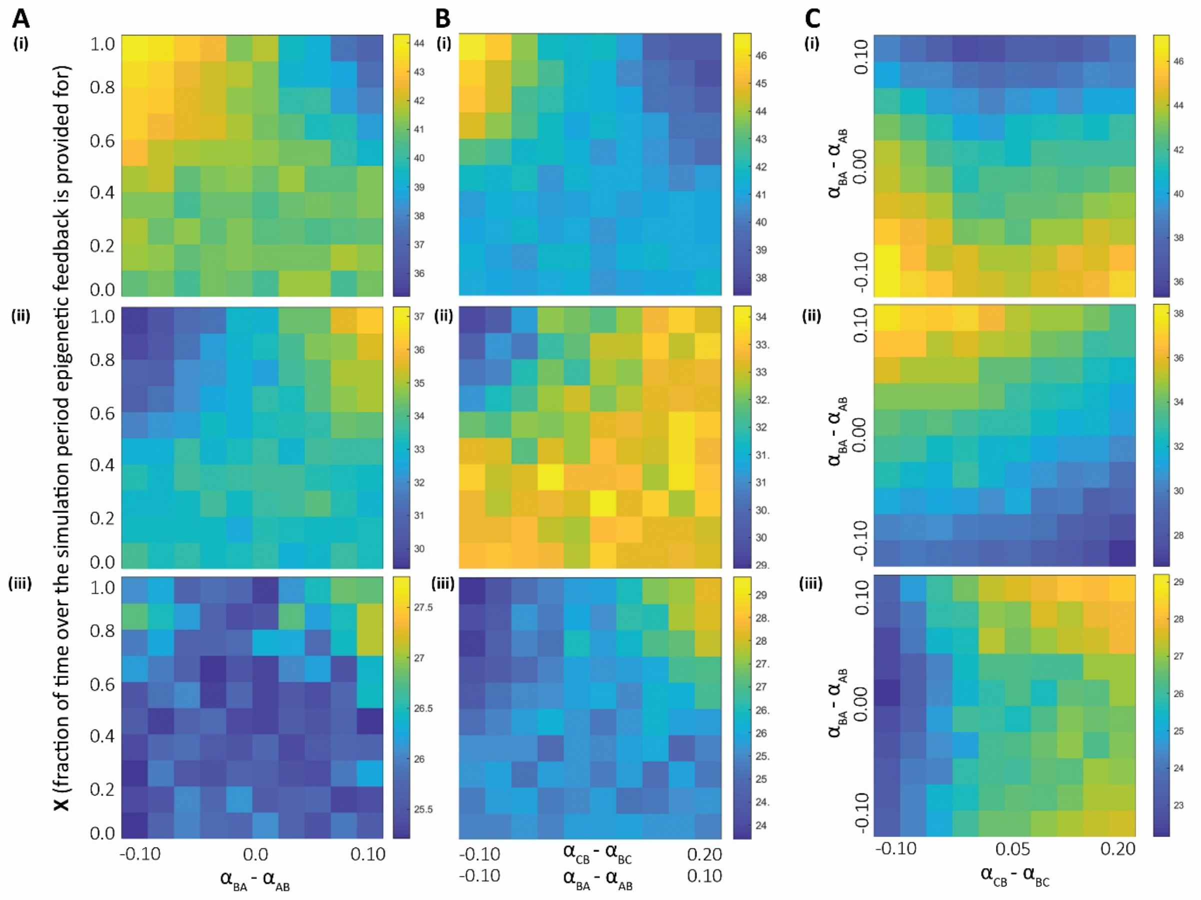


***Fig S16: A) i)*** *Phase plot of population percentage corresponding to node A with variations in two parameters: X and the relative epigenetic influence which is varied from a case of stronger repression from A to B ($\alpha_{BA} < \alpha_{AB}$) to a stronger repression from B to A ($\alpha_{BA} > \alpha_{AB}$)* ***ii)*** *Same as* ***i)*** *but for state B.* ***iii)*** *Same as* ***i)*** *but for state C.* ***B) i)*** *Same as A;* ***i)*** *but for varying relative epigenetic influence in both feedback loops (between A and B, and between C and B).* ***ii)*** *Same as* ***i)*** *but for state B.* ***iii)*** *Same as* ***i)*** *but for state C.* ***C) i)*** *Phase plot of population percentage corresponding to state A with bifurcation parameters as the difference in α values corresponding to mutual epigenetic repression ($\alpha_{BA}$ - $\alpha_{AB}$ and $\alpha_{CB}$ - $\alpha_{BC}$), at X = 1.* ***ii)*** *Same as* ***i)*** *but for state B.* ***iii)*** *Same as* ***i)*** *but for state C. Parameter set P3 (Table S1) is used here.*

### Fig S17

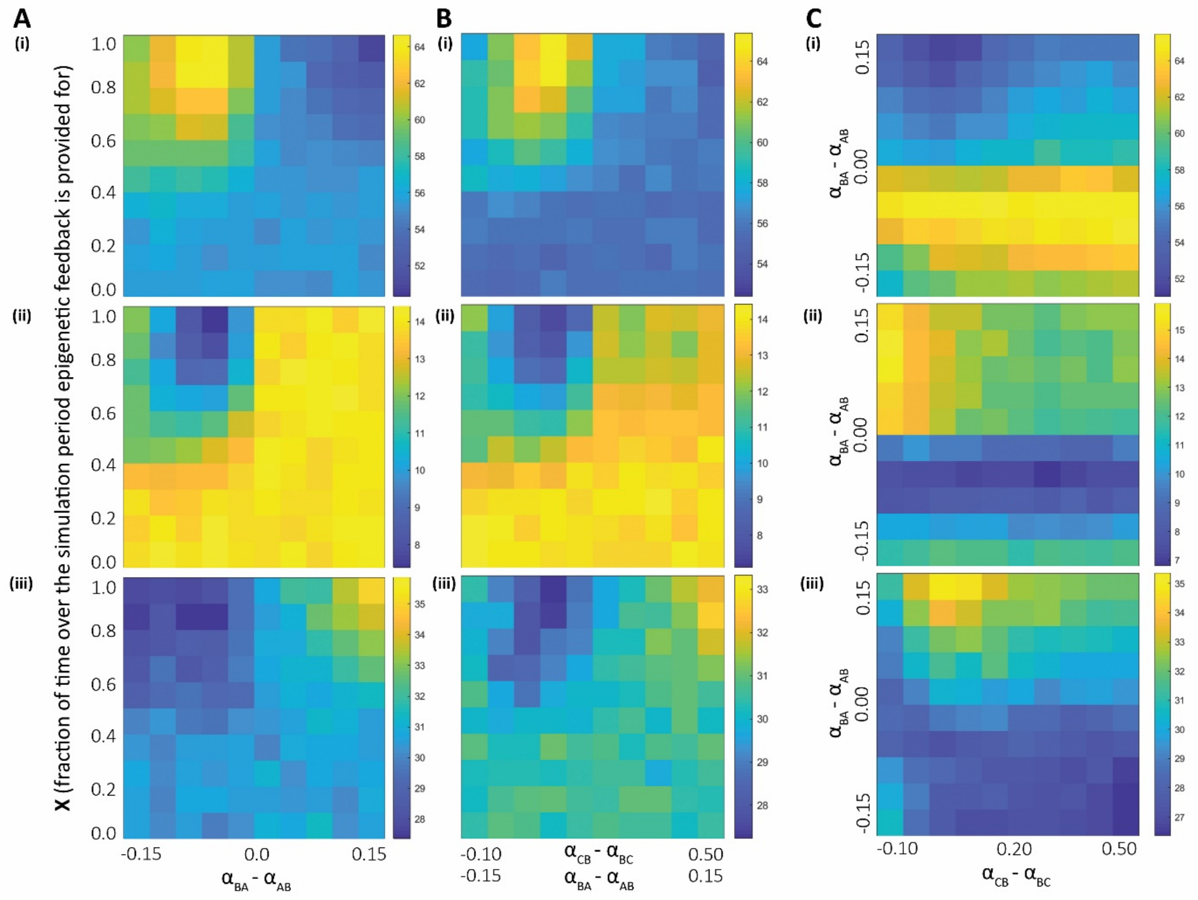


***Fig S17: A)i)*** *Phase plot of population percentage corresponding to node A with variations in two parameters: X and the relative epigenetic influence which is varied from a case of stronger repression from A to B ($\alpha_{BA} < \alpha_{AB}$) to a stronger repression from B to A ($\alpha_{BA} > \alpha_{AB}$)* ***ii)*** *Same as* ***i)*** *but for state B.* ***iii)*** *Same as* ***i)*** *but for state C.* ***B)i)*** *Same as A;* ***i)*** *but for varying relative epigenetic influence in both feedback loops (between A and B, and between C and B).* ***ii)*** *Same as* ***i)*** *but for state B.* ***iii)*** *Same as* ***i)*** *but for state C.* ***C) i)*** *Phase plot of population percentage corresponding to state A with bifurcation parameters as the difference in α values corresponding to mutual epigenetic repression ($\alpha_{BA}$ - $\alpha_{AB}$ and $\alpha_{CB}$ - $\alpha_{BC}$), at X = 1.* ***ii)*** *Same as* ***i)*** *but for state B.* ***iii)*** *Same as* ***i)*** *but for state C. Parameter set P4 (Table S1) is used here.*

### Fig S18

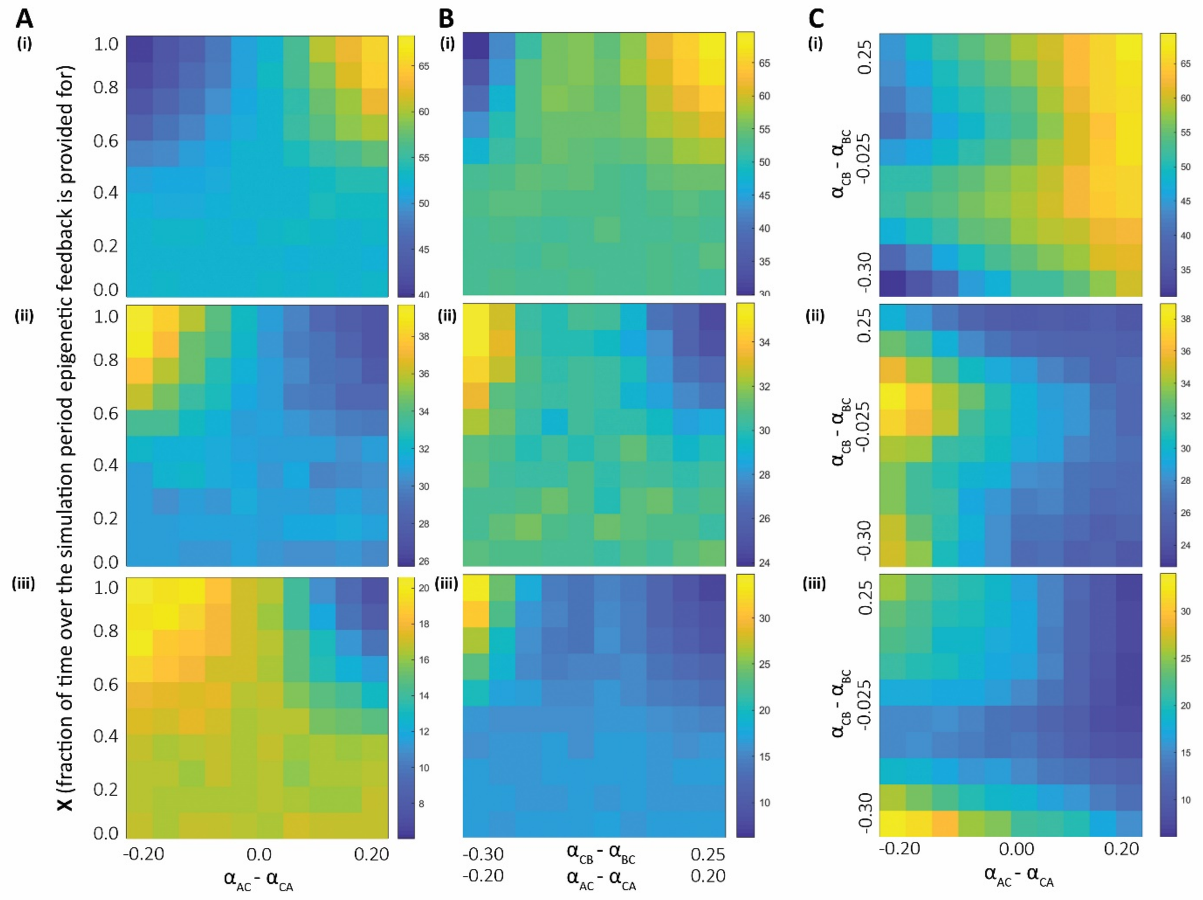


***Fig S18: A) i)** Phase plot of population percentage corresponding to node A with variations in two parameters: X and the relative epigenetic influence which is varied from a case of stronger repression from C to A ($\alpha_{AC} < \alpha_{CA}$) to a stronger repression from A to C ($\alpha_{AC} > \alpha_{CA}$) **ii)** Same as **i)** but for state B. **iii)** Same as **i)** but for state C. **B) i)** Same as A; **i)** but for varying relative epigenetic influence in both feedback loops (between A and C, and between B and C). **ii)** Same as **i)** but for state B. **iii)** Same as **i)** but for state C. **C) i)** Phase plot of population percentage corresponding to state A with bifurcation parameters as the difference in α values corresponding to mutual epigenetic repression ($\alpha_{CB}$ - $\alpha_{BC}$ and $\alpha_{AC}$ - $\alpha_{CA}$), at X = 1. **ii)** Same as **i)** but for state B. **iii)** Same as **i)** but for state C. Parameter set P5 (Table S1) is used here.*
